# Dendritic cell Piezo1 integrating mechanical stiffness and inflammatory signals directs the differentiation of T_H_1 and T_reg_ cells in cancer

**DOI:** 10.1101/2022.05.17.492270

**Authors:** Yuexin Wang, Hui Yang, Anna Jia, Yufei Wang, Qiuli Yang, Yingjie Dong, Yujing Bi, Guangwei Liu

**Affiliations:** Key Laboratory of Cell Proliferation and Regulation Biology, Ministry of Education, College of Life Sciences, Beijing Normal University, Beijing 100875 China; Department of Immunology, School of Basic Medical Sciences, Fudan University, Shanghai 200032 China; State Key Laboratory of Pathogen and Biosecurity, Beijing Institute of Microbiology and Epidemiology, Beijing 100071 China

**Keywords:** dendritic cells, Piezo1, innate immunity, T cell differentiation, T_H_1 cells, T_reg_ cells, glycolysis, HIF1α, cancer, tumor microenvironment

## Abstract

Dendritic cells (DCs) play an important role in anti-tumor immunity by inducing T cell differentiation. Herein, we found that the mechanical sensor Piezo1 expressed by DCs integrates innate inflammatory stimuli and stiffness signals and directs the reciprocal differentiation of T_H_1 and regulatory T (T_reg_) cells in cancer. Genetic deletion of Piezo1 in DCs inhibited the generation of T_H_1 cells while driving the development of T_reg_ cells in promoting cancer growth. Mechanistically, Piezo1-deficient DCs regulated the secretion of the polarizing cytokines TGFβ1 and IL-12, leading to increased TGFβR2-p-Smad3 activity and decreased IL-12Rβ2-p-STAT4 activity while inducing the reciprocal differentiation of T_reg_ and T_H_1 cells. In addition, Piezo1 integrated the SIRT1-hypoxia-inducible factor-1 alpha (HIF1α)-dependent metabolic pathway and calcium-calcineurin-NFAT signaling pathway to orchestrate reciprocal T_H_1 and T_reg_ lineage commitment through DC-derived IL-12 and TGFβ1. Our studies provide critical insight for understanding the role of the DC-based mechanical regulation of immunopathology in directing T cell lineage commitment in tumor microenvironments.

## Introduction

CD4^+^ T helper cells often play a central role in cancer, through their differentiation into different T cell subsets(1, 2). At present, many studies have shown that naïve CD4^+^ T cells differentiate into many kinds of effector cells, including T_H_1 cells, T_H_2 cells, T_H_9 cells, T_H_17 cells and regulatory T (T_reg_) cells, in specific cytokine environments(3–5). In tumor microenvironment, in addition to the regulatory role of CD4^+^ T cells and the underlying mechanisms, foreign antigens and all kinds of innate stimuli are often presented by antigen-presenting cells (APCs) to further direct the development and differentiation of different subsets of CD4^+^ T cells(6, 7). Innate inflammatory stimuli, such as bacterial lipopolysaccharide (LPS), cytokine or physiological stimuli, such as oxygen, nutrient availability, and even force and pressure, can also change the immune response. In particular, the tumor microenvironment integrates different innate physiological or pathological stimuli and develop a complex stimulation microenvironment (8, 9). Response to various exogenous and endogenous stimuli, DCs, as professional APCs, bridge innate and adaptive and regulate the differentiation of different T cell subsets largely through three key signaling pathways, including costimulatory molecule expression, T cell receptor (TCR) signaling and cytokine production(10–17). DC-derived cytokines and chemokines exert proinflammatory or anti-inflammatory effects and are involved in the shaping of distinct T cell subset lineage programs to determine the prognosis of cancer patients. For example, DCs can secrete polarizing cytokines, such as TGFβ1 and IL-12. TGFβ1 can induce the production of Foxp3^+^ T_reg_ cells by activating TGFβ receptor (TGFβR) on the surface of T cells, while IL-12 can stimulate IL-12R on the surface of T cells to induce the differentiation of IFNγ^+^ T_H_1 cells. Studies from our lab and others have shown that innate signaling in DCs or myeloid-derived cells mediated by S1P1, Hippo kinase MST1, mitogen-activated protein kinase (MAPKs), calcineurin-NFAT and Wnt-β-catenin plays a critical role in programming naïve CD4^+^ T cell differentiation(18–23). How the differentiation of CD4^+^ T cells is modulated and regulated by innate immune signaling pathways in DCs in the tumor microenvironment remains unclear.

Piezo1 was originally identified as a mechanically activated non-selective cation ion channel with significant permeability to calcium ions, is evolutionally conserved and is involved in the proliferation and development of various types of cells in the context of various types of mechanical and/or innate stimuli. It has been reported that innate inflammatory stimulation or mechanical changes, such as changes in stiffness, can activate Piezo1 and trigger an inflammatory response(4, 9). Piezo1 exerts significant regulatory effects on many kinds of immune cells. It has been reported that Piezo1 can regulate the functions of macrophages, DCs and T cells in inflammation and cancer(8, 9, 24–27). However, it is still unclear whether Piezo1-targeted DCs affect the differentiation of different subsets of T cells in cancer.

Here, we found that the mechanical sensor Piezo1 integrates innate inflammatory stimuli and stiffness signals and directs the reciprocal differentiation of T_H_1 and regulatory T (T_reg_) cells in inhibiting tumor growth. Piezo1 largely acts by integrating SIRT1-hypoxia-inducible factor-1 alpha (HIF1α)-dependent metabolic activity and calcium-calcineurin-nuclear factor of activated T cells (NFAT) signaling, and regulates the production of polarizing cytokines IL-12 and TGF-β1 by DCs as well as the expression of IL-12Rβ2 and TGFβR2 by responding T cells.

## Results

### Inflammatory and stiffness stimuli alter Piezo1 expressions by DCs

As previously reported(26, 28), environmental stiffness can alter the secretion of inflammatory factors by DCs. Moreover, the mechanical force receptor Piezo1 may be involved in this regulation. We first examined the effect of environmental stiffness on the expression of piezo1 in DCs. To investigate the effect of substrate stiffness on the function of DCs, we used a cell culture system consisting of a poly two methyl siloxane (PDMS) hydrogel-coated plate, which was previously described(4, 28). The mechanical properties of the PDMS hydrogel can be changed by adjusting the matrix/curing agent ratio, and this ratio can be precisely adjusted to simulate physiological tension. Consistent with previous reports(4, 28), we used 2 kPa to mimic lymphoid tissue under physiological conditions and 50 kPa to mimic lymphoid tissue under inflammatory conditions. Sorted splenic DCs cultured on a stiff hydrogel (E=50 kPa) or plastic plates exhibited significantly enhanced expression of the proinflammatory cytokine IL-12p40 and diminished expression of the anti-inflammatory cytokine TGFβ1 compared with DCs cultured on a pliant hydrogel (E=2 kPa; Fig. S1A). This suggests that substrate stiffness could regulate inflammatory cytokine secretion by DCs. Moreover, the expression of the mechanical force receptor Piezo1 was significantly upregulated in 50 kPa-conditioned hydrogels compared with 2 kPa-conditioned hydrogels, similar to that in DCs stimulated with the inflammatory stimulus LPS (Fig. S1B). Although 50 kPa-conditioned hydrogels significantly caused more IL-12p40 and less TGFβ1 compared with 2 kPa, mechanical force receptor Piezo1 deficient DC cells significant rescue them to normal level (Fig. S1C). These data suggested that mechanical force receptor Piezo1 mediated the inflammatory cytokine production induced by substrate stiffness in DCs.

### DC-specific Piezo1-deficient mice exhibited altered T cell differentiation

To investigate the regulatory effect of piezo1 expressed by DCs on T cell function, we generated DC-specific piezo1 conditional knockout mice with *Piezo1*^flox/flox^ and *Cd11c-cre*, and these mice were called *Piezo1*^ΔDC^ mice. *Piezo1*^ΔDC^ mice showed no obvious abnormalities after birth. However, after 40 weeks of age, *Piezo1*^ΔDC^ mice showed lower weight loss than WT mice (Fig. S2A). Moreover, *Piezo1*^ΔDC^ mice showed reduced T cell activation in mesenteric lymph nodes (MLNs), peyer’s patches (PPs), intraepithelial lymphocytes (IELs) and lamina propria lymphocytes (LPLs) (Fig. S2B-C). Importantly, Piezo1 deficiency in DCs results in fewer IFNγ^+^ T_H_1 cells and more Foxp3^+^ T_reg_ cells but does not affect the numbers of T_H_2 or T_H_17 cells (Fig. S3A-B). Thus, we conclude that DC-specific Piezo1 deficiency alters the differentiation of T_H_1 and T_reg_ cells during aged-related T cell responses.

### DC-specific Piezo1 regulates T cell differentiation in promoting tumor growth

Next, we studied the effect of DC-specific piezo1 deletion on T cell differentiation in MC38 mouse colon cancer. We observed changes in tumor growth in *Piezo1*^ΔDC^ and WT mice. The rate of tumor growth was significantly faster and greater in *Piezo1*^ΔDC^ than in WT mice (Fig. 1A). Flow cytometry showed that *Piezo1*^ΔDC^ mice had more Foxp3^+^ T_reg_ cells, fewer IFNγ^+^ T_H_1 cells and normal numbers of T_H_2 and T_H_17 cells in tumor tissue compared with WT control groups (Fig. 1B). Furthermore, to observe the dynamic process of T cell differentiation during the tumor environment, we isolated T cells from tumor-bearing mice at day 20, 30, and 40 and observed dynamic T cell differentiation. Both T-bet and IFNγ levels were downregulated rapidly, Foxp3 expression in T cells was upregulated gradually, and IL-4, IL-17A, and GATA3 expression did not change in *Piezo1*^ΔDC^ mice compared with WT mice (Fig. 1C). Thus, these data suggest that DC-specific Piezo1 deficiency probably directs T_reg_ and T_H_1 cell differentiation to promote the tumor growth in the context of tumor microenvironment.

**Figure 1.**
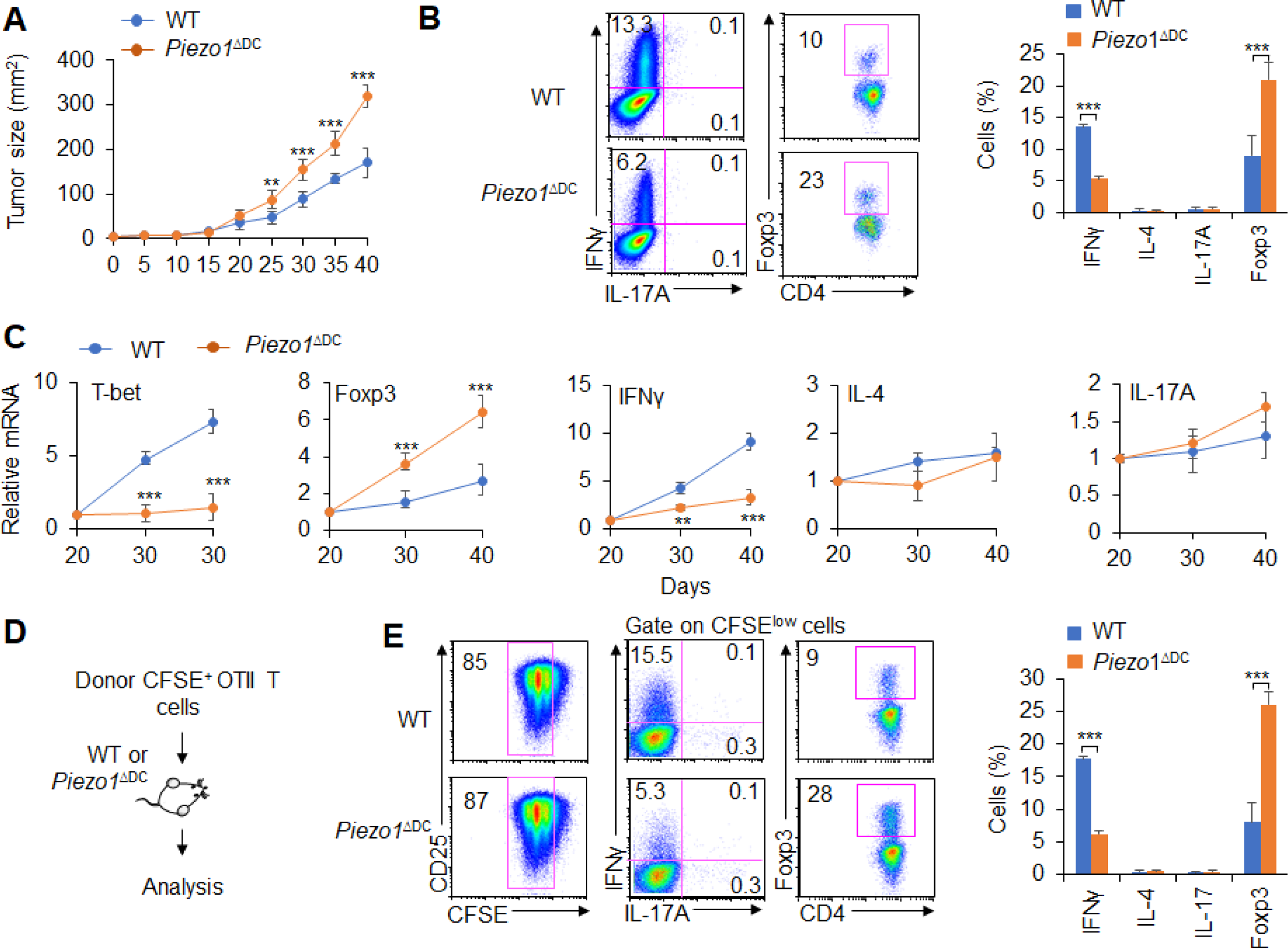
DC-specific Piezo1 regulates T cell differentiation in cancer. (**A**) MC38 tumor cells were implanted subcutaneously in WT and *Piezo1*^ΔDC^ mice (n=10) and tumor size was measured every 5 days for 40 days. (**B**) Intracellular staining of IFNγ, IL-4, IL-17A and Foxp3 expression by CD4^+^ T cells sorting from the tumor of WT and *Piezo1*^ΔDC^ tumor-bearing mice at Day 40. (**C**) mRNA expression of the indicated genes by CD4^+^ T cells isolated from the draining lymph nodes (dLNs) of tumor from WT and *Piezo1*^ΔDC^ tumor-bearing mice on the indicated days (the levels in WT mice at Day 20 were set to 1). (**D**) MC38 tumor cells were implanted subcutaneously in WT and *Piezo1*^ΔDC^ mice (n=10) and at Day 20, the CD45.1^+^ donor CFSE^+^OT II CD4^+^T cells were transferred into WT and *Piezo1*^ΔDC^ tumor-bearing mice for 10 days. The CD45.1^+^ CFSE^+^ donor T cells were analyzed and the intracellular staining of IFNγ, IL-4, IL-17A and Foxp3 expression among CFSE^low^ donor T cells. The data are representative of three to four independent experiments (mean ± s.d.; n=4). \*\**P*<0.01, and \*\*\**P*<0.001, compared with the indicated groups.

Further, we observed the antigen specific responses of T cells in tumor-bearing mice during tumor growth. MC38-OVA tumor cells were implanted subcutaneously in WT and *Piezo1*^ΔDC^ mice. And at Day 20 after tumor cell implantation, naïve CD45.1^+^ OTII T cells were isolated from OT II-TCR transgenic mice, labeled with CFSE and transferred into WT and *Piezo1*^ΔDC^ bearing-tumor recipient mice (Fig. 1D). After 10 days of adoptive transfer of OTII T cells to recipient tumor-bearing mice, the intracellular staining of the CD45.1^+^ CFSE^+^ donor T cells from tumors in recipient WT and *Piezo1*^ΔDC^ mice were analyzed showed that the percentage of CFSE^+^ T cells is similar between WT and *Piezo1*^ΔDC^ bearing tumor mice. However, there are more Foxp3^+^T_reg_ cells and less IFNγ^+^ T_H_1 cells, but similar T_H_2 and T_H_17 cells in WT and *Piezo1*^ΔDC^ bearing-tumor mice (Fig. 1D-E). These data suggest that Piezo1 in DCs regulates the T_H_1 and T_reg_ differentiation in tumor microenvironment with the antigen-specific manner.

### DC-specific Piezo1 expression instructs antigen-specific T_H_1 and T_reg_ differentiation

Next, we conducted an adoptive transfer experiment to investigate the T cell response induced by DCs expressing Piezo1. Naïve T cells (CD45.2^+^CD4^+^TCR^+^CD44^low^ CD62L^high^) from C57BL/6 mice were transferred into CD45.1^+^C57BL/6 recipient mice, and then, the recipient mice were immunized with WT and *Piezo1*^ΔDC^ splenic DCs and LPS. The donor cells were analyzed on Day 10 after immunization. T cell proliferation was comparable between the WT and *Piezo1*^ΔDC^ mice (Fig. 2A). However, the donor mice immunized with *Piezo1*^ΔDC^ splenic DCs exhibited a higher percentage of Foxp3^+^CD4^+^ T cells and a lower percentage of IFNγ^+^CD4^+^ T cells than WT control mice. Both the WT and *Piezo1*^ΔDC^ mice showed similar levels of IL-4 and IL-17 expression among donor CD4^+^ T cells (Fig. 2B-C).

**Figure 2.**
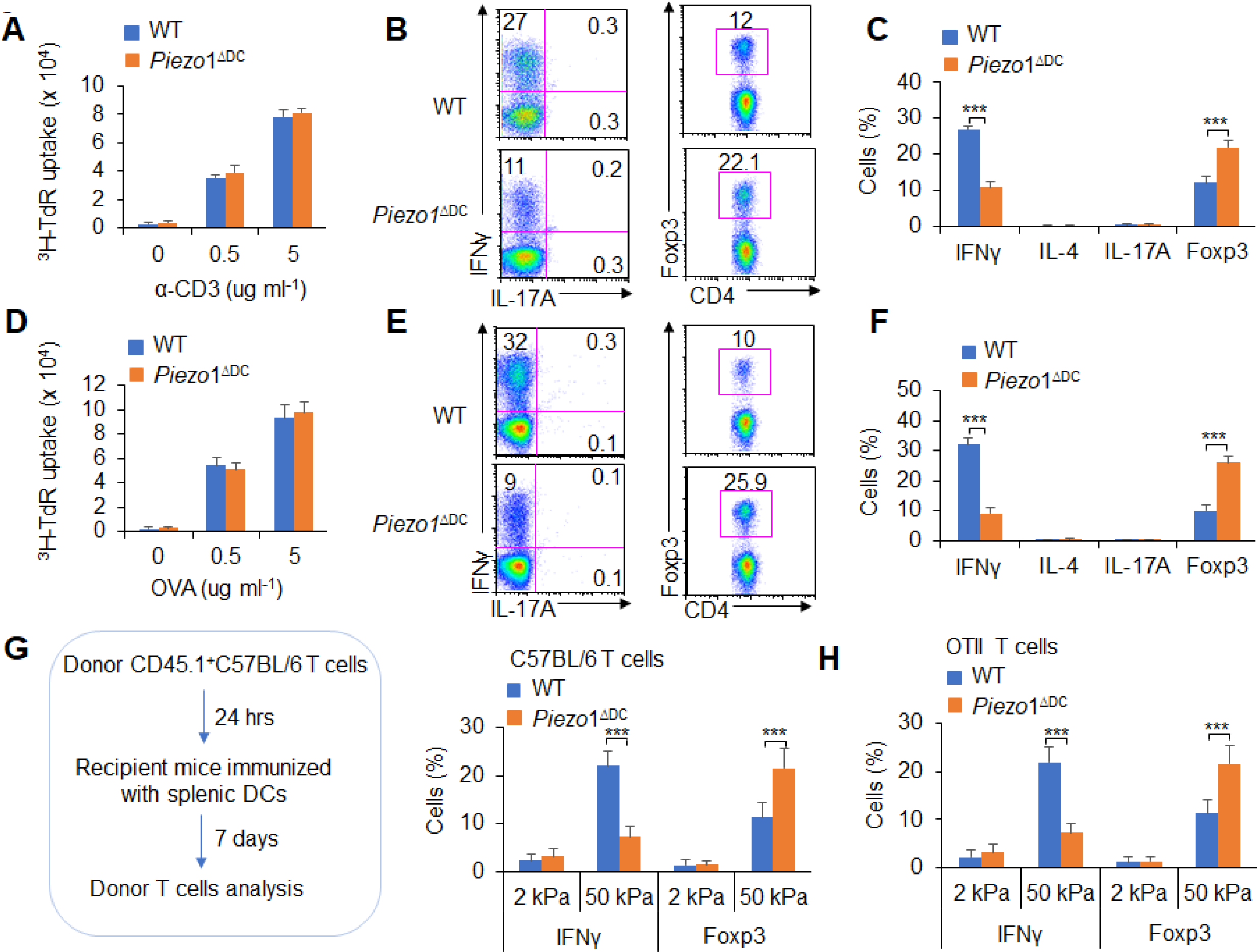
DC-specific Piezo1 expression directs the differentiation of T_H_1 and T_reg_ cells in vivo. (**A-C**) Naïve CD45.2^+^ T cells were transferred into CD45.1^+^ C57BL/6 WT mice, and the mice were immunized with WT and *Piezo1*^ΔDC^ splenic DCs and LPS. DLN cells were analyzed at Day 7 after immunization. (A) Donor CD45.2^+^ T cell proliferation after stimulation with anti-CD3 (2 µg/ml) and anti-CD28 (2 μg/ml) antibodies. (B-C) Intracellular staining of IFNγ, IL-4, IL-17A and Foxp3 expression in donor-derived (CD45.2^+^) CD4^+^ T cells after PMA and ionomycin stimulation. A representative figure is shown in B, and the data are summarized in C. (**D-F**) Naïve OT-II T cells were transferred into CD45.1^+^ C57BL/6 WT mice, and the mice were immunized with WT and *Piezo1*^ΔDC^ splenic DCs and OVA+CFA. DLN cells were analyzed at Day 7 after immunization. (D) Donor CD45.2^+^ T cell proliferation after stimulation with OVA. (E-F) Intracellular staining of IFNγ, IL-4, IL-17A and Foxp3 in donor-derived (CD45.2^+^) CD4^+^ T cells after OVA stimulation. A representative figure is shown in E, and the data are summarized in F. (**G-H**) Splenic DCs isolated from WT and *Piezo1*^ΔDC^ mice were plated on 2 kPa and 50 kPa hydrogels and incubated for 24 hrs. (G) Naïve T cells were transferred into CD45.1^+^ C57BL/6 WT mice, and the mice were immunized with 2 kPa and 50 kPa hydrogel-conditioned DCs. DLN cells were analyzed at Day 7 after immunization. Intracellular staining of IFNγ, IL-4 and IL-17A in donor-derived (CD45.2^+^) CD4^+^ T cells after PMA and ionomycin stimulation. A representative figure is shown on the left, and the data are summarized on the right. (H) Naïve OT-II T cells were transferred into CD45.1^+^ C57BL/6 WT mice, and the mice were immunized with 2 kPa or 50 kPa hydrogel-conditioned DCs and OVA+CFA. DLN cells were analyzed at Day 7 after immunization. Intracellular staining of IFNγ, IL-4 and IL-17A in donor-derived (CD45.2^+^) CD4^+^ T cells after OVA stimulation. A representative figure is shown on the left, and the data are summarized on the right. The data are representative of three to four independent experiments (mean ± s.d.; n=3-4). \*\*\**P*<0.001, compared with the indicated groups.

Furthermore, we observed the antigen-specific response of adoptive transfer donor T cells whose differentiation was induced by DC-specific Piezo1 expression. Naïve OTII T cells were isolated and transferred into CD45.1^+^ C57BL/6 recipient mice, and then, the recipient mice were immunized with WT and *Piezo1*^ΔDC^ DCs and OVA in the presence of complete Freund’s adjuvant (CFA). The donor cells were analyzed on Day 10 after immunization. When T cells isolated from the draining lymph node (DLN) were stimulated with OVA antigen, WT and *Piezo1*^ΔDC^ showed similar T cell proliferation effects (Fig. 2D). However, donor cells immunized with *Piezo1*^ΔDC^ DCs exhibited more Foxp3^+^ cells and fewer IFNγ^+^ cells than the WT control cells. Both WT and *Piezo1*^ΔDC^ mice showed similar numbers of IL-4^+^ T_H_2 cells and IL-17A^+^ T_H_17 cells (Fig. 2E-F). Together, these data showed that Piezo1 deficiency in DCs induced antigen-specific T cell differentiation in vivo.

To further observe the T cell response induced by DCs exposed to stiffness stimuli, we further examined DCs conditioned by 2 kPa or 50 kPa hydrogel in adoptive transfer experiments. Splenic DCs were isolated from WT and *Piezo1*^ΔDC^ mice and plated on 2 kPa or 50 kPa hydrogels. Naïve T cells were isolated and transferred into CD45.1^+^ recipient mice, and then, the recipient mice were immunized with 2 kPa or 50 kPa hydrogel-conditioned DCs. Donor T cells from mice immunized with *Piezo1*^ΔDC^ DCs conditioned with 50 kPa, but not 2 kPa hydrogels included more Foxp3^+^ T_reg_ cells and less IFNγ^+^ T_H_1 cells (Fig. 2G). Similarly, naïve OTII T cells were transferred into recipient mice, which were immunized with WT or *Piezo1*^ΔDC^ DCs conditioned by 2 kPa or 50 kPa hydrogel and OVA in the presence of CFA. The donor cells from mice immunized with *Piezo1*^ΔDC^ DCs conditioned with 50 kPa, but not 2 kPa hydrogels included more Foxp3^+^ T_reg_ cells and fewer IFNγ^+^ T_H_1 cells (Fig. 2H). Thus, the mechanical sensor Piezo1 in DCs could integrate innate inflammatory and stiffness stimuli and direct the reciprocal differentiation of T_H_1 and T_reg_ cells.

### Piezo1 is required for DC-dependent T cell differentiation

Next, we investigated the effects of Piezo1 expression by DCs on T cell subset differentiation in an in vitro system. Polyclonal T cells cocultured with *Piezo1*^ΔDC^ DCs in the presence of anti-CD3 and LPS induced more Foxp3^+^T_reg_ cell and less IFNγ^+^ T_H_1 cell, even with a variety of *Piezo1*^ΔDC^ DCs including splenic CD11b^+^ DC or CD8α^+^ DCs, than WT control DCs (Fig. 3A and Fig. S4A). Furthermore, antigen-specific OTII T cells cocultured with *Piezo1*^ΔDC^ splenic DCs, including the CD11b^+^ DC and CD8^+^ DC subsets, in the presence of antigen (OVA) and LPS induced more Foxp3^+^ T_reg_ cells and fewer IFNγ^+^ T_H_1 cells (Fig. 3B and Fig. S4B). Moreover, T-bet and IFNγ expression was significantly downregulated and Foxp3 expression was significantly upregulated in T cells cocultured with *Piezo1*^ΔDC^ DCs compared with those cocultured with WT compartment control DCs. However, IL-4 and IL-17A expression was similar in T cells cocultured with *Piezo1*^ΔDC^ DCs or WT DCs (Fig. 3C). These data suggest that Piezo1 deficiency in different kinds of DCs including CD11b^+^ DC and CD8^+^ DC subsets regulates the reciprocal differentiation of T_H_1 and T_reg_ cells in an antigen-specific manner.

**Figure 3.**
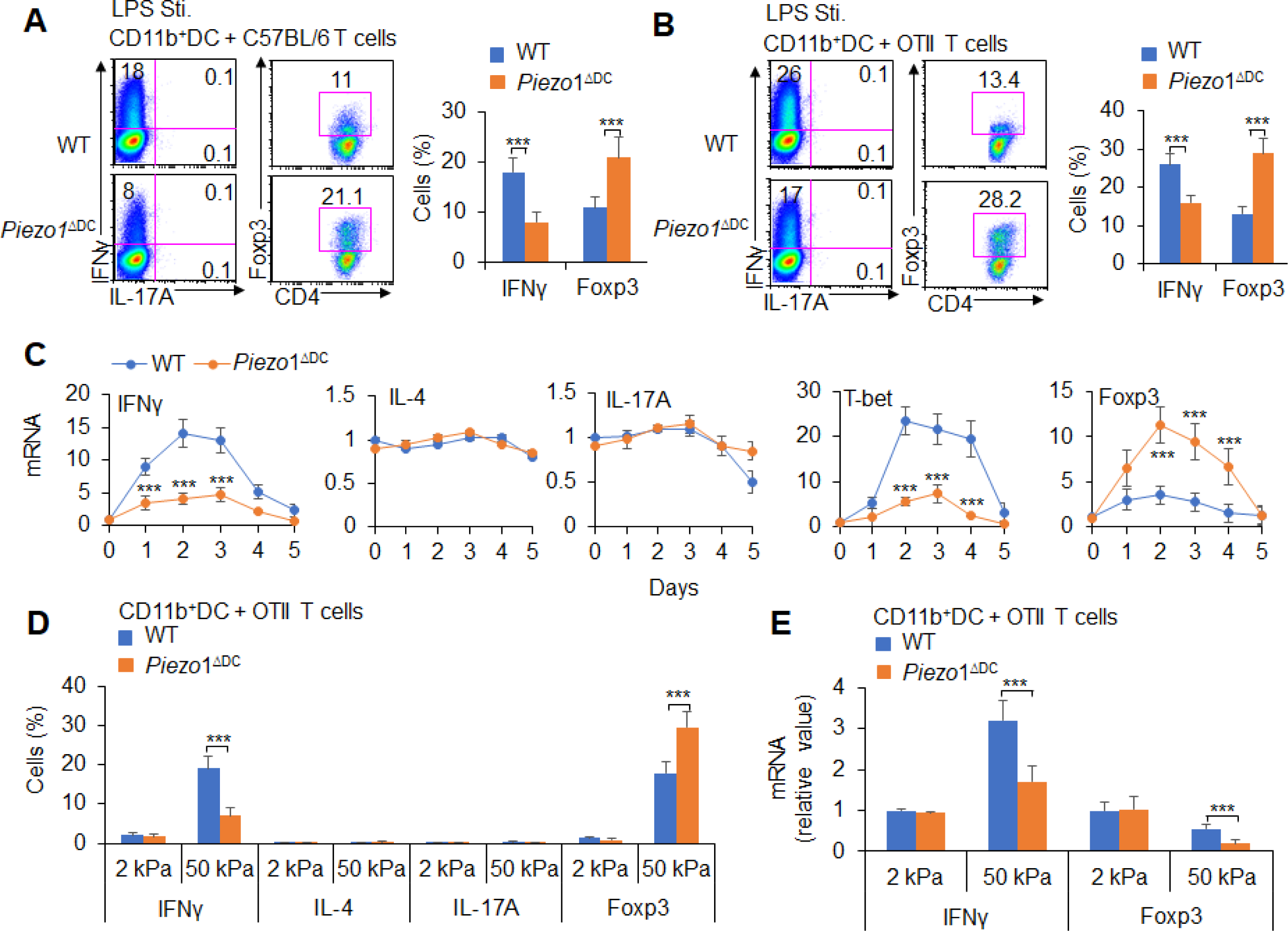
DC-specific Piezo1 expression directs T_H_1 and T_reg_ differentiation in vitro. (**A**) C57BL/6 naïve CD4^+^ T cells were cocultured with LPS-pulsed splenic CD11b^+^ DCs from WT and *Piezo1*^ΔDC^ mice for 5 days. Intracellular staining of IFNγ and Foxp3 in T cells. A representative figure is shown on the left, and the data are summarized on the right. (**B**) Naïve CD4^+^ T cells from OT II mice were cocultured with LPS-pulsed splenic CD11b^+^ DCs from WT and *Piezo1*^ΔDC^ mice for 5 days. Intracellular staining of IFNγ and Foxp3 in T cells. A representative figure is shown on the left, and the data are summarized on the right. (**C**) Naïve T cells from C57BL/6 mice were cocultured with splenic DCs from WT and *Piezo1*^ΔDC^ mice for 5 days in the presence of anti-CD3 (2 ng/ml) and LPS (10 ng/ml). The relative mRNA expression of the indicated genes in T cells was determined with qPCR. The levels in the WT group and on Day 0 were set to 1. (**D**) splenic CD11b^+^ DCs isolated from WT and *Piezo1*^ΔDC^ mice were plated on 2 kPa and 50 kPa hydrogels (1 x 10^5^ cells/well) for 24 hrs. Naïve CD4^+^ T cells from OT II mice were cocultured with 2 kPa and 50 kPa hydrogel-conditioned DCs in the presence of OVA (5 μg/ml) for 5 days. Intracellular staining of IFNγ, IL-4, IL-17A and Foxp3 in T cells. A representative figure is shown on the left, and the data are summarized on the right. (**E**) Naïve T cells from OTII mice were cocultured with 2 kPa and 50 kPa hydrogel-conditioned splenic CD11b^+^ DCs in the presence of OVA (5 μg/ml) for 5 days. The relative mRNA expression of the indicated genes in T cells was determined with qPCR. The levels in the WT group with 2 kPa were set to 1. The data are representative of three to four independent experiments (mean ± s.d.; n=3-6). \*\*\**P*<0.001, compared with the indicated groups.

In addition to inflammatory LPS stimuli, we also observed the response to stiffness stimuli. Coculture of antigen-specific OTII T cells with *Piezo1*^ΔDC^ splenic CD11b^+^ DCs conditioned by 2 kPa or 50 kPa hydrogels in the presence of antigen induced a higher percentage and mRNA expression of Foxp3 and a lower percentage and mRNA expression of IFNγ than coculture of antigen-specific OTII T cells with WT control DCs (Fig. 3D-E). Together, Piezo1 signals from DCs integrating innate inflammatory and stiffness stimuli directs reciprocal T_H_1 and T_reg_ cell differentiation in an antigen-specific manner.

### DC Piezo1 regulates T cell differentiation through IL-12 and TGFβ1

APCs regulate T cell differentiation by changing costimulatory molecule expression, TCR signaling and polarizing cytokine production(6). The expression of MHC and the costimulatory molecules CD80, CD86 and CD54 was comparable between WT and *Piezo1*^ΔDC^ DCs (Fig. S5). We also detected changes in cytokines secreted by splenic DCs, especially polarizing cytokines important for inducing T_H_1 and T_reg_ cell differentiation. We found that *Piezo1^Δ^*^DC^ caused significantly higher TGFβ1 expression and lower IL-12p40 expression than WT control DCs in the presence of LPS (10 ng/ml; Fig. 4A) or 50 kPa, but not 2 kPa hydrogels (Fig. 4B). These results suggest that the polarizing cytokines TGFβ1 and IL-12 are probably involved in regulating T cell subset differentiation induced by Piezo1 in DCs stimulating by inflammatory stimuli or stiffness signals.

**Figure 4.**
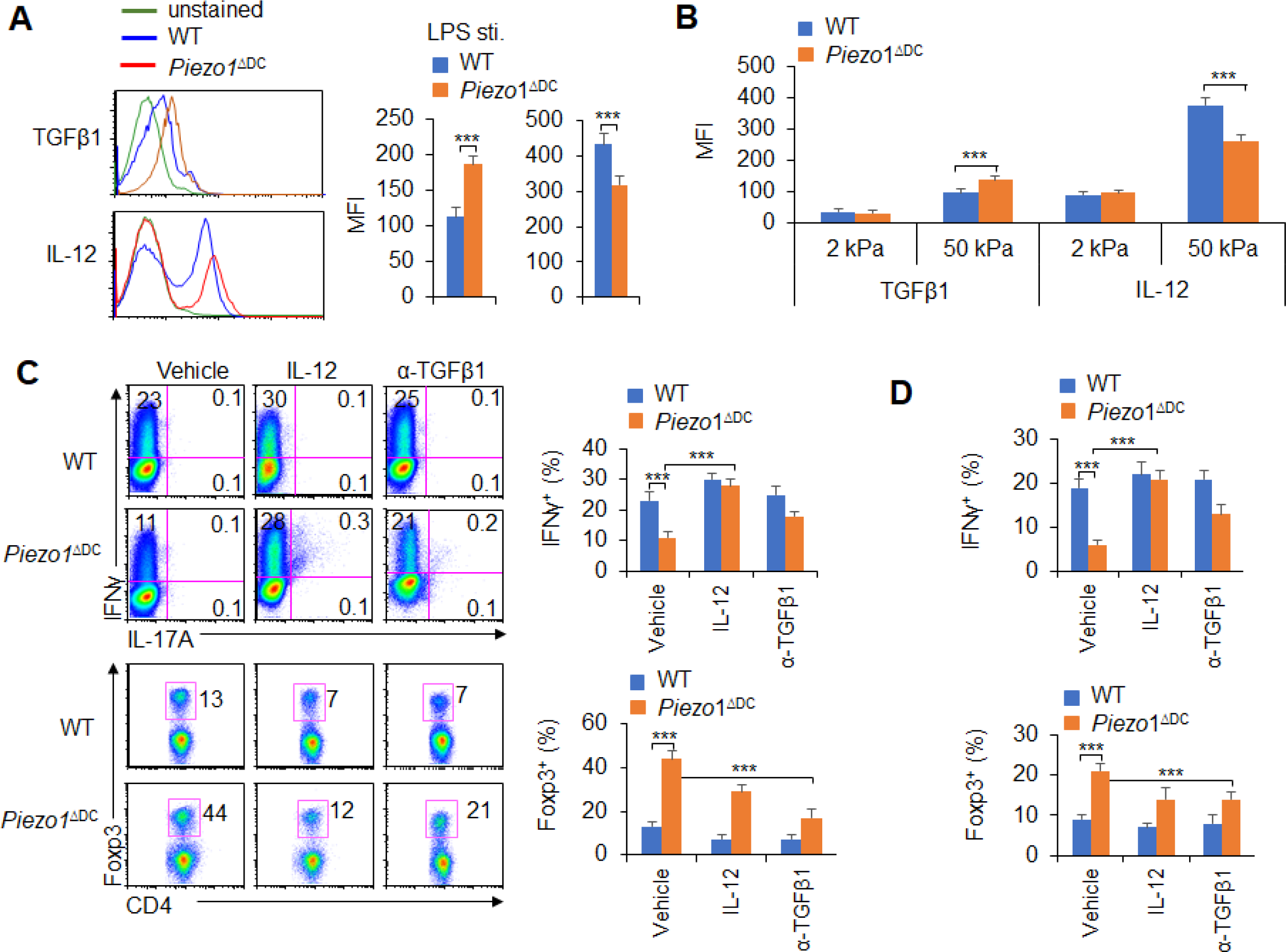
Piezo1 regulates IL-12 and TGFβ1 production by DCs to direct T_H_1 and T_reg_ cell differentiation. (**A-B**) Intracellular staining of IL-12p40 and TGFβ1 expression in WT and *Piezo1*^ΔDC^ splenic DCs after 5 hrs of treatment with LPS (A; 10 ng/ml) or culture on 2 kPa and 50 kPa hydrogels (B). A representative figure is shown on the left, and the data are summarized on the right. (**C**) Intracellular staining of IFNγ and Foxp3 in T cells cocultured with WT and *Piezo1*^ΔDC^ splenic DCs in the presence of the indicated treatments (IL-12, Peprotech, 10 μg/ml or anti-TGFβ1, R&D Systems, 20 μg/ml) for 5 days. A representative figure is shown on the left, and the data are summarized on the right. (**D**) Intracellular staining of IFNγ (upper panel) and Foxp3 (lower panel) in T cells cocultured with WT and *Piezo1*^ΔDC^ splenic DCs conditioned with 50 kPa hydrogel and the indicated treatments for 5 days. Data are summarized. The data are representative of three to four independent experiments (mean ± s.d.; n=3-5). \*\*\**P*<0.001, compared with the indicated groups.

We selected a splenic DC-T coculture system to determine whether DC-specific Piezo1 expression regulates T cell subset differentiation through the polarizing cytokines TGFβ1 and IL-12. Although Piezo1 deficiency in DCs caused a significantly lower IFNγ^+^ T_H_1 cell percentage, adding IL-12 to the coculture system almost completely recovered the proportion of *Piezo1*^ΔDC^ DCs during T_H_1 cell differentiation (Fig. 4C). Even DCs conditioned by 50 kPa hydrogel treatment caused similar effects (Fig. 4D). Consistently, blocking TGFβ1 signaling with an anti-TGFβ1 antibody in the coculture system almost completely recovered the effect of *Piezo1*^ΔDC^ splenic DCs on T_reg_ cell differentiation. Similar effects could be observed when DCs were treated with LPS or 50 kPa hydrogels (Fig. 4C-D). Thus, we could conclude that DC-specific Piezo1 expression regulates the reciprocal differentiation of T_H_1 and T_reg_ cells through the polarizing cytokines IL-12 and TGFβ1.

### DC-specific Piezo1 expression induces T cell differentiation through TGFβR2 and IL-12Rβ2

T cell differentiation-inducing cytokines often change the corresponding receptors on the surface of T cells and program the differentiation of T cell subsets(1, 19). We further determined the corresponding receptor expression on T cells. Piezo1 deficiency in splenic DCs significantly increased the expression of TGFβR2 and decreased the expression of IL-12Rβ2 but did not affect the expression of TGFβR1/3 or IL-12Rβ1 on T cells in a DC-T coculture system (Fig. 5A and Fig. S6A-B). Interestingly, Piezo1-deficient DCs also exhibited significantly higher levels of phosphorylated Smad3 and lower levels of phosphorylated STAT4 than WT DCs (Fig. 5B). These data suggest that TGFβR2-p-Smad3 or IL-12Rβ2-p-STAT4 signaling is involved in the T_reg_ and T_H_1 cell differentiation induced by DC-specific Piezo1 expression.

**Figure 5.**
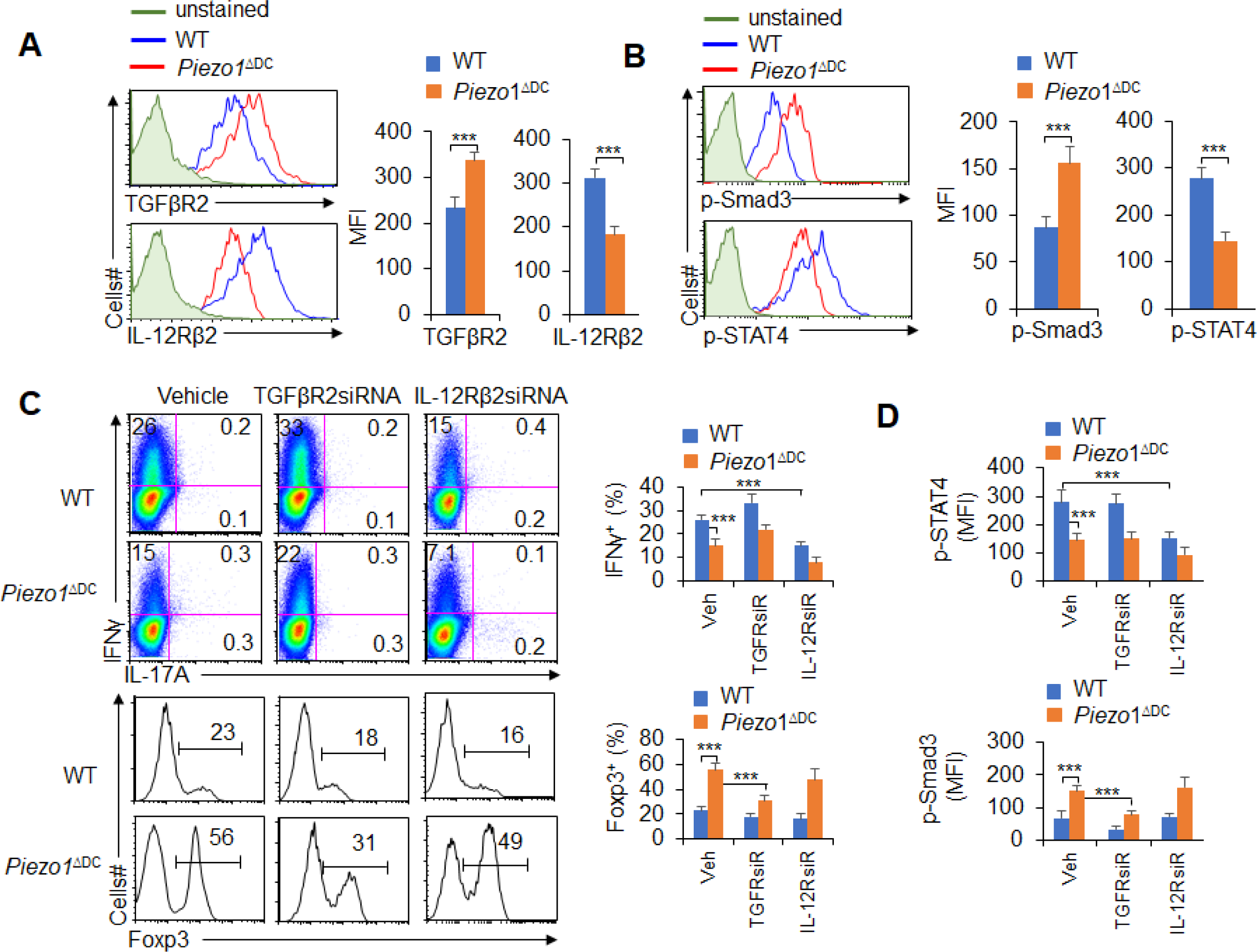
TGFβR2-Smad3 and IL-12Rβ2-pSTAT4 are required for the T cell differentiation induced by DC-specific Piezo1 expression. (**A**) Expression of TGFβR2 and IL-12Rβ2 in T cells cocultured with WT or *Piezo1*^ΔDC^ splenic DCs for 5 days. A representative figure is shown on the left, and the data are summarized on the right. (**B**) Intracellular staining of p-Smad3 and p-STAT4 in T cells cocultured with WT or *Piezo1*^ΔDC^ splenic DCs for 5 days. A representative figure is shown on the left, and the data are summarized on the right. (**C-D**) Sorted naïve T cells were transfected with control, TGFβR2 shRNA vector or IL-12Rβ2 shRNA vector and cocultured with WT or *Piezo1*^ΔDC^ DCs for 5 days. Intracellular staining of IFNγ (C; upper panel) and Foxp3 (C; lower panel) in T cells. A representative figure is shown on the left, and the data are summarized on the right. (D) Intracellular staining of p-Smad3 and p-STAT4 in T cells. Data are summarized. The data are representative of three to four independent experiments (mean ± s.d.; n=4). \*\*\**P*<0.001, compared with the indicated groups.

To determine whether TGFβR2-pSmda3 and IL-12Rβ2-pSTAT4 are required for promoting the T cell differentiation induced by DC-specific Piezo1 expression, we knocked down IL-12Rβ2 and TGFβR2 expression in T cells with shRNA in a DC-T coculture system (Fig. S7A-B). Although DC-specific Piezo1 deficiency resulted in significantly more Foxp3^+^ T_reg_ cells, higher phosphorylation of Smad3, less IFNγ^+^ T_H_1 cells, and lower phosphorylation of STAT4, knockdown of TGFβR2 expression significantly recovered the effects compared with the WT conditions (Fig. 5C-D). Consistently, knockdown of IL-12Rβ2 expression significantly decreased the percentage of IFNγ^+^ T_H_1 cells and the expression of p-STAT4 in the WT and *Piezo1*^ΔDC^ groups (Fig. 5C-D). These data suggest that TGFβR2-pSmda3 and IL-12Rβ2-pSTAT4 signaling in T cells are required for the T cell differentiation induced by DC Piezo1.

### Piezo1 regulates IL-12 and TGFβ1 production through the SIRT1-HIF1α-glycolysis pathway

How does Piezo1 regulate IL-12 and TGFβ1 production to direct T cell differentiation? To study the mechanisms underlying the effects of Piezo1, splenic DCs were stimulated by LPS, and we assessed the signaling downstream of LPS stimulation, including Erk, c-jun-NH2-kinase (JNK), p38, SIRT1, HIF1α and glycolytic molecular signaling.

Glucose can be used to fuel ATP production through two linked metabolic pathways: glycolysis and oxidative phosphorylation (OXPHOS; including the tricarboxylic acid cycle [TCA]) during immune responses. We investigated the role of glycolysis and OXPHOS signal activities in the functional regulation of DCs induced by Piezo1. It showed that LPS treatment led to an increase in the proton production rate (PPR), but splenic DCs treated with the Piezo1 agonist Yoda1 exhibited significantly enhanced PPR values but not oxygen consumption rates (OCRs) (Fig. S8A-B). Consistently, Yoda1 treatment significantly upregulated the expression of glycolytic molecules in splenic DCs (Fig. S8C). Thus, these data suggest glycolysis, but not OXPHOS signal pathway, probably involved in the DC functional regulation induced by Piezo1. Blocking glycolysis with 2-deoxy-D-glucose (2-DG), a prototypical inhibitor of glycolysis pathways, significantly recovered the IL-12 and TGFβ1 production in splenic DCs induced by Yoda1 treatment (Fig. S9A-B). Furthermore, Piezo1 deficiency in DCs significantly decreased the PPR value and the expression of glycolytic molecules but not the OCR value (Fig. 6A-C and Fig. S10A). Blocking glycolysis with 2-DG significantly recovered the productions of IL-12 and TGFβ1 in *Piezo1*^ΔDC^ DCs to normal level compared with WT DCs (Fig. 6A-C and Fig. S10B-C). These data suggest that glycolysis activities are required for the polarizing cytokine production in DCs induced by Piezo1.

**Figure 6.**
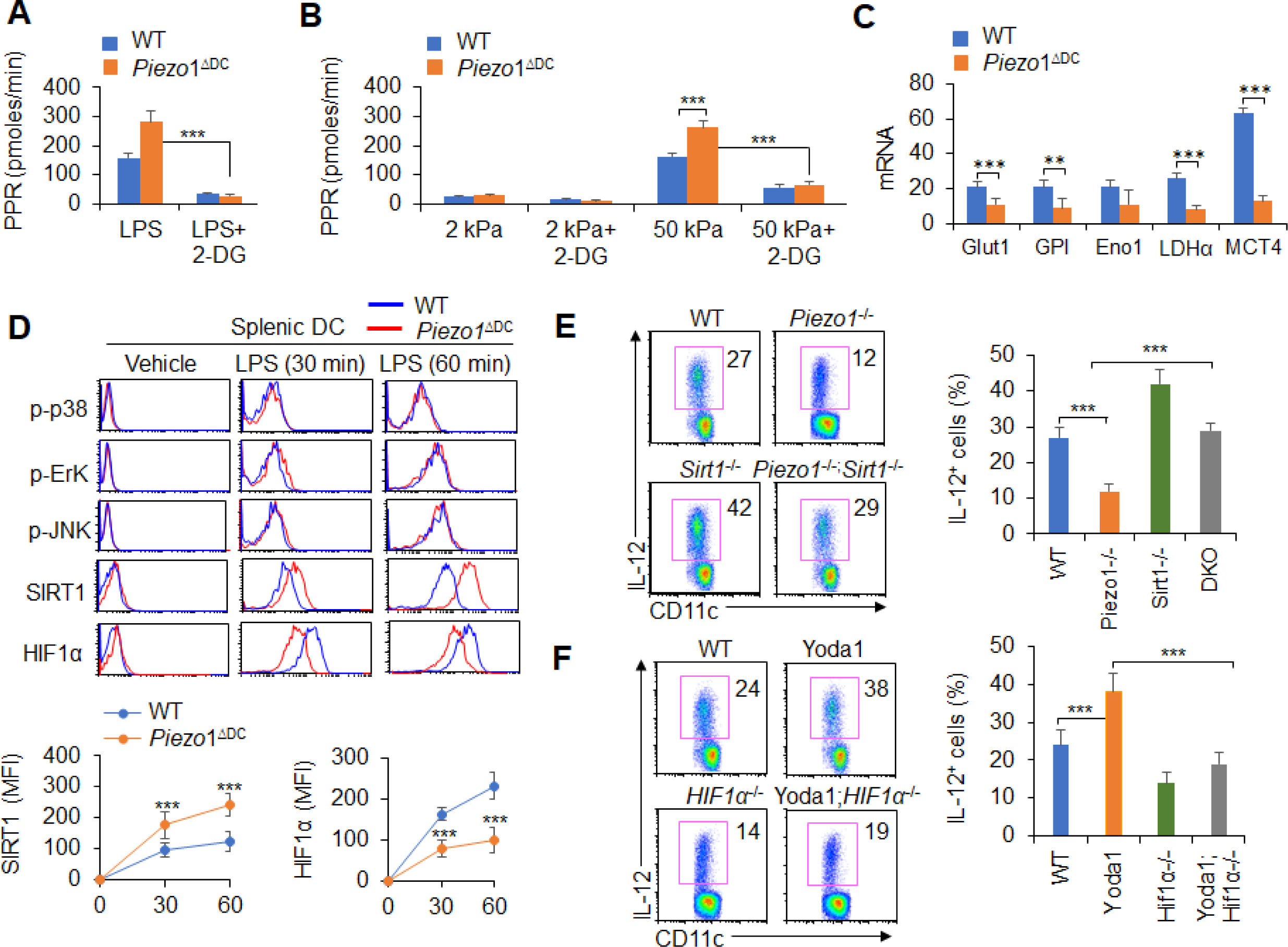
Piezo1 regulates TGFβ1 and IL-12 production through the SIRT1-HIF1α-glycolysis pathway. (**A-B**) Sorted splenic DCs from WT or *Piezo1*^ΔDC^ mice were stimulated with LPS (10 ng/ml; A) or with 2 kPa or 50 kPa hydrogels (B) for 24 hrs in the presence or absence of 2-DG (1 mmol/l). The PPR was analyzed as a readout for glycolysis. (**C**) mRNA expression of glycolytic molecules in splenic DCs from WT or *Piezo1*^ΔDC^ mice treated with LPS (10 ng/ml) for 12 hrs. The levels in the WT control group were set to 1. (**D**) Intracellular staining of p38, Erk, and JNK phosphorylation and SIRT1 and HIF1α expression in splenic DCs from WT or *Piezo1*^ΔDC^ mice. A representative figure is shown in the upper panel, and the data are summarized in the lower panel. (**E**) Splenic DCs from WT, *Piezo1*^ΔDC^ (*Piezo1*^-/-^), *Sirt1*^-/-^, and *Piezo1*/*Sirt1* double knockout (DKO; *Piezo1*^-/-^*Sirt1*^-/-^) mice were stimulated with LPS (10 ng/ml). Intracellular staining of IL-12p40. A representative figure is shown on the left, and the data are summarized on the right. (**F**) Splenic DCs from WT or *Hif1α*^-/-^ mice were stimulated with LPS (10 ng/ml) in the presence or absence of Yoda1 (25 μM). Intracellular staining of IL-12p40. A representative figure is shown on the left, and the data are summarized on the right. The data are representative of three to four independent experiments (mean ± s.d.; n=3-4). \*\**P*<0.01 and \*\*\**P*<0.001, compared with the indicated groups.

Additionally, as expected, LPS activated all downstream pathways in WT DCs including p-JNK, p-Erk, p-p38, SIRT1 and HIF1α signal pathway. However, Piezo1 deletion in DCs enhanced the phosphorylation of Erk, p38 and JNK, similar to WT DCs. However, *Piezo1*^ΔDC^ DCs exhibited stronger and sustained activation of the histone deacetylase SIRT1 and weaker and shorter activation of the transcription factor HIF1α (Fig. 6D). Thus, Piezo1 is probably associated with SIRT1-HIF1α signaling and glycolysis activation.

To determine whether SIRT1 is involved in this regulation, we used Piezo1 and SIRT1 double knockout (DKO) mice in this investigation. Interestingly, less IL-12 and more TGFβ1 production in *Piezo1*^ΔDC^ DCs was significantly reversed in Piezo1-SIRT1 DKO cells (Fig. 6E and Fig. S11A). Consistently, HIF1α expression and glycolysis activities were significantly recovered to a normal level (Fig. S11B-C). These data suggest SIRT1 is required for the IL-12 and TGFβ1 production in DCs induced by Piezo1 and HIF1α and glycolysis activation is probably related with these alterations.

To determine whether HIF1α is involved in this regulation, we crossed DC HIF1α conditional knockout mice (*Hif1α*^ΔDC^) with *Hif1α*^flox/flox^ and *Cd11c-Cre* mice. Splenic DCs were isolated from WT and *Hif1α*^ΔDC^ mice and treated with the Piezo1 agonist Yoda1. More IL-12 and less TGFβ1 in Yoda1 treated DCs was significantly reversed in Yoda1 treated *Hif1α*^ΔDC^ DCs (Fig. 6F and Fig. S12A). These data suggest HIF1α is required for the IL-12 and TGFβ1 production in DCs induced by Piezo1.

Moreover, splenic DCs were treated with the Piezo1 agonist Yoda1, which significantly altered glycolysis activity and SIRT1 expression in DCs. SIRT1 expressions cannot be altered in *Hif1α*^ΔDC^ DCs compared with control group (Fig. S12B). However, treatment of *Hif1α*^ΔDC^ with Yoda1 significantly reversed the alteration of glycolysis but not SIRT1 expression (Fig. S12B-C). These suggest HIF1α and glycolysis signals are probably the downstream targets of SIRT1 in regulating cytokine production in DCs induced by Piezo1. Therefore, these data collectively suggest that HIF1α and glycolysis is downstream targets of SIRT1 in regulating the Piezo1-induced cytokine production in DCs.

### Piezo1 regulates IL-12 and TGFβ1 production through the calcium-calcineurin-NFAT axis

Piezo1 is a Ca^2+^-dependent mechanosensitive channel(9). PIEZO1 was recently shown to mediate both antigen receptor-driven and stretch- or cyclical force-driven mechanical sensors, and it plays an important role in macrophage mechanosensory signal transduction and regulates DC function(25, 26). Therefore, we assessed the level of calcium influx in *Piezo1*^ΔDC^ splenic DCs. Both inflammatory stimulation and stiffness stimulation caused a significant decrease in the calcium influx in *Piezo1*^ΔDC^ DCs compared with the WT control cells (Fig. 7A). And, Piezo1 agonist Yoda1 treatment significantly enhanced intracellular calcium influx, and also decreased TGFβ1 secretion and increased IL-12 secretion by DCs exposed to LPS or conditioned by 50 kPa hydrogels (Fig. 7B-D). Importantly, blocking the Ca^2+^ signaling pathway with ruthenium red reversed these alterations. However, another nonspecific ion channel inhibitor, gadolinium, had no effects on calcium influx or cytokine production in DCs (Fig. 7B-D). These data collectively suggest calcium influx is required for the cytokine production in DCs induced by Piezo1. Consistently, *Piezo1*^ΔDC^ DCs showed lower calcium influx and exhibited higher TGFβ1 and lower IL-12 levels, and blocking the Ca^2+^ signaling pathway with ruthenium red, but not gadolinium reversed these alterations in cytokine production (Fig. S13A-C). Thus, calcium signaling pathway is required for the TGFβ1 and IL-12 production in DCs induced by Piezo1.

**Figure 7.**
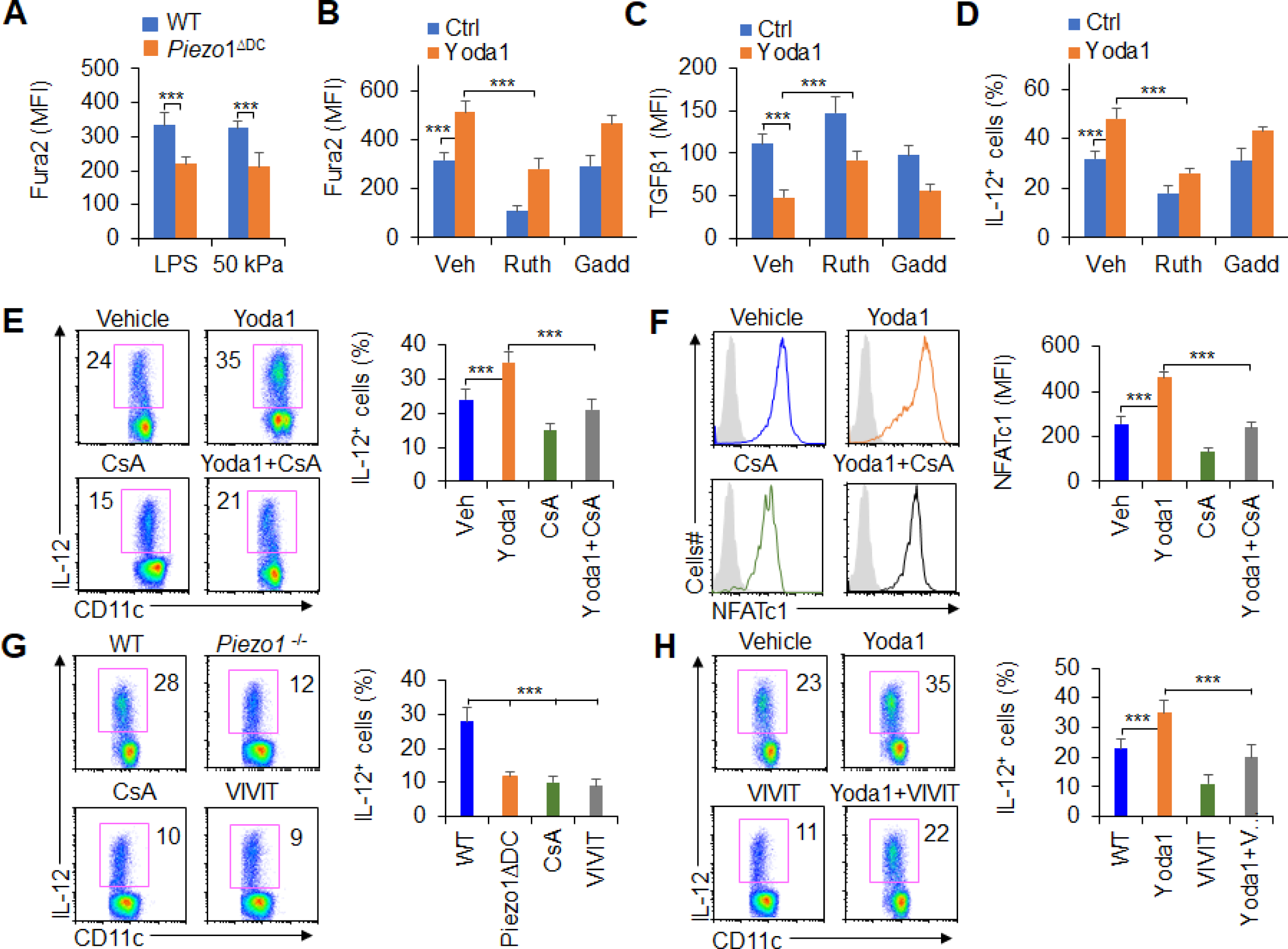
Piezo1 regulates TGFβ1 and IL-12 production through the calcium-calcineurin-NFAT axis. (**A**) Measurement of intracellular Ca^2+^ concentrations with Fura2 dye in splenic DCs from WT or *Piezo1*^ΔDC^ mice treated with LPS (10 ng/ml) or cultured on plates containing 50 kPa hydrogels. (**B**) Intracellular Ca^2+^ concentrations measured with Fura2 in splenic DCs from WT mice after the indicated treatment (Yoda1, 25 μM, MCE; ruthenium red, 30 μM, Sigma; gadolinium chloride, 10 μM, Sigma). (**C-D**) Intracellular staining of TGFβ1 (C) and IL-12p40 (D) in splenic DCs from WT mice after the indicated treatments. (**E**) Intracellular staining of IL-12p40 in splenic DCs from WT mice after the indicated treatments. A representative figure is shown on the left, and the data are summarized on the right. (**F**) Intracellular staining of NFATc1 in splenic DCs from WT mice after the indicated treatments (CsA, 10 nM). A representative figure is shown on the left, and the data are summarized on the right. (**G-H**) Intracellular staining of IL-12p40 in splenic DCs from WT or *Piezo1*^ΔDC^ mice after the indicated treatments (Yoda1, 25 μM, MCE; 11R-VIVIT, 100 nM, MCE; CsA, 10 nM, Sigma). A representative figure is shown on the left, and the data are summarized on the right. The data are representative of three to four independent experiments (mean ± s.d.; n=3-4). \*\*\**P*<0.001, compared with the indicated groups.

Our previous studies and other studies have shown that calcineurin-NFAT are critical molecules of the calcium signaling pathway in regulating the immune response(22, 29–31). These prompted us to investigate whether calcineurin-NFAT signals are necessary for Piezo1 to regulate DC function through calcium signaling pathway. Therefore, we pharmacologically targeted calcineurin and NFAT to assess their roles in regulating Piezo1-induced TGFβ1 and IL-12 production in DCs. Splenic DCs treated with the Piezo1 agonist Yoda1 caused more IL-12 and less TGFβ1 production, but blocking calcineurin with its inhibitor cyclosporin A (CsA) reversed these alterations in DCs (Fig. 7E and Fig. S14A). These results suggest that calcium-calcineurin signaling are probably required for cytokine productions in DCs induced by Piezo1.

As reported previously(22, 29–31), NFAT is critical transactional factor for regulating calcium-calcineurin signaling pathway in mediating immune cell activities. Moreover, upregulation of Piezo1 with Yoda1 treatment significantly enhanced the expression of NFAT, blocking calcineurin with CsA significantly inhibits the expression of NFAT (Fig. 7F). To test the role of NFAT for Piezo1 to regulate DC function, we pharmacologically targeted NFAT with its inhibitor VIVT to assess the role of NFAT in regulating Piezo1-induced TGFβ1 and IL-12 production by DCs. *Piezo1*^ΔDC^ DCs exhibited lower IL-12 and higher TGFβ1 production, and blocking calcineurin with CsA and blocking NFAT with VIVIT consistently showed similar cytokine production by DCs (Fig. 7G and Fig. S14B). Interestingly, splenic DCs treated with the Piezo1 agonist Yoda1 exhibited more IL-12 and less TGFβ1 production, but blocking NFAT with VIVIT reversed these alterations in DCs (Fig. 7H and Fig. S14C). Altogether, these data suggest that the calcium-calcineurin-NFAT axis is necessary for regulating IL-12 and TGFβ1 production by DC Piezo1.

## Discussion

DCs play a central role in initiating first-line innate immunity and inducing subsequent adaptive immunity in protecting against tumorigenesis (32, 33). As a professional APCs, DCs can efficiently shape antigen-specific adaptive immune responses by presenting various exogenous and endogenous antigen stimuli, regulating cell surface costimulatory molecule expression, and producing cytokines and chemokines(34, 35). Fine-tuning a myriad of DC intrinsic signaling pathways is required to induce an effective adaptive immune response without triggering inflammation-induced host injury(35, 36). Innate inflammatory stimuli include inflammatory stimuli, oxygen, nutrient availability, and even force and pressure, often change the DC responses and affect the immune outcome in diseases. Especially, tumor microenvironment usually integrates different innate inflammatory and stiffness stimuli and develop a complex stimulation microenvironment, but how does DC integrate inflammatory and stiffness stimuli and regulates T cell differentiation in tumor remains unclear. Here, our data revealed that the mechanical sensor Piezo1, a signal node, responds to innate inflammatory and stiffness stimuli and integrates both the SIRT1-HIF1α-glycolysis metabolic signaling axis and calcium-calcineurin-NFAT signaling in DCs to drive T_H_1 differentiation while inhibiting T_reg_ lineage commitment in inhibiting tumor growth in the context of complex tumor microenvironment. The changes in IL-12Rβ2/TGFβR2 expression and downstream STAT4/SMAD3 signaling in responding T cells further result in strong DC-T cell crosstalk, indicating the differentiation of T_reg_ and T_H_1 cells (Fig. S15). Thus, our results can contribute to a more comprehensive understanding of the immunopathological process of DC Piezo1-derived T cell differentiation in the tumor microenvironment.

Recent studies have suggested that Piezo1 is involved in regulating many of diseases, including the infectious inflammation and cancer(37–41). Piezo1 modulates macrophage polarization and stiffness sensing, which are related to calcium influx and the promotion of macrophage activation by actin(25). The Piezo1-mediated response to LPS inflammatory stimulation mediates cell activation through TLR4 and then mediates the function of macrophages in digesting and killing pathogens through calcium influx and the Camkii-Mst1/2-Rac signaling pathway(9). Global inhibition with a peptide inhibitor showed protective effects against both cancer and septic shock, which are associated with decreased numbers of myeloid-derived suppressor cells (MDSCs)(24). In addition to inflammatory stimulation, Piezo1 also showed immune regulatory effects on mechanical stimulation. Cyclical hydrostatic pressure initiates an inflammatory response via the mechanically activated ion channel Piezo1. Piezo1 deficiency in innate immune cells ablated pulmonary inflammation in the context of bacterial infection or fibrotic autoinflammation(8). Additionally, mechanical stiffness controls DC metabolism and function(26). Although these studies have clearly shown that Piezo1 can respond to inflammatory stimulation as well as induce an immune response to mechanical stimulation and regulate immune cell functions, especially in innate immune cells, it is still unclear how to target innate APCs, especially DCs, to direct T cell differentiation in cancer. Piezo1, a mechanically activated ion channel with high affinity for calcium, is sensitive to both innate inflammatory stimuli and mechanical stiffness and more easily triggers the calcium signaling pathway to mediate the T cell immune response. Our results showed that Piezo1 could respond to both inflammatory stimulation and stiffness, and subsequently, Piezo1 could effectively integrate metabolism signals and ion signals pathway including SIRT1-HIF1α-glycolysis and calcium influx and the calcium-calcineurin-NFAT signaling pathway to direct the differentiation of T_reg_ and T_H_1 cells by regulating the production of DC-derived polarizing cytokines, including IL-12 and TGFβ1, in the context of tumor microenvironment (Fig. S15).

As a transcription factor, HIF1α has been implicated as a critical proinflammatory signaling module in myeloid leukocytes(42–47). Consistent with recent findings that SIRT1 is responsible for the deacetylation and destabilization of HIF1α(48, 49). HIF1α is critically involved in regulating Piezo1-induced innate immune cell function(8). The metabolic mechanism is probably critical in regulating DC function(26). HIF1α-dependent glycolysis metabolism is also critical for regulating Th9 differentiation and MDSC development and function(50). The present data showed that Piezo1 could target SIRT1-HIF1α-glycolysis metabolism signaling to modulate DC-derived polarizing cytokine secretion. As an ion channel, Piezo1 is sensitive to calcium influx in regulating immune cell function (25). These data also showed that blocking calcium influx significantly altered Piezo1-mediated DC-derived cytokine secretion. Previous studies have shown that calcineurin-NFAT in MDSCs regulates T_reg_ function(22). Here, we further showed that Piezo1 targeting the calcineurin-NFAT axis modulates DC-derived polarizing cytokine secretion to direct T_reg_ and T_H_1 cell differentiation in cancer.

Metabolic regulation and cellular signals are closely and generally associated with the immune response, but there are still few studies on the regulation of the calcium signaling pathway and immune response(51–53). Our data showed that Piezo1 could integrate the metabolic signaling and calcium signaling pathways to modulate DC-derived polarizing cytokine secretion in the context of cancerous inflammation. Effective immune responses require DCs to function under various conditions, including altered extracellular mechanical tension states, intracellular metabolic states, and ion levels (possibly caused by inflammatory stimulation) or due to migration due to nutritional and/or hypoxic environments (tumor microenvironment). The adaptation of DCs to changing metabolic states and calcium signaling results from a mechanism of a “mechanical sensor checkpoint”, an active signaling process involved in sensing changes in metabolic and intracellular calcium levels and subsequent signaling transduction and execution(54–57). Our data further suggested that the mechanical sensor Piezo1 in DCs requires the interplay of metabolic and intracellular calcium checkpoints, including the metabolic SIRT1-HIF1α-glycolysis pathway and a sensitive calcium signaling pathway, calcium-calcineurin-NFAT signaling(58, 59).

In summary, targeting the mechanical sensor Piezo1 in DCs alters the secretion of the polarizing cytokines TGFβ1 and IL-12 and the expression of TGFβR2-pSmad3 and IL-12Rβ2-pSTAT4, thereby contributing to the reciprocal differentiation of T_reg_ and T_H_1 cells in promoting cancer growth. Thus, our results define the essential nature of Piezo1 as a signaling node that confers innate inflammatory and stiffness signals to initiate a cooperation between the Piezo1-SIRT1-HIF1α-glycolysis metabolism signaling pathway and calcium-Piezo1-calcineurin-NFAT ion signaling pathway in DCs. This signaling induces the reciprocal differentiation of T_reg_ cells and T_H_1 cells and has implications for targeting DCs as an approach to the treatment of cancer.

## Materials and Methods

### Mice

All the animal experiments were performed with the approval of the Animal Ethics Committee of Fudan University, China and Beijing Normal University, China. C57BL/6 *Piezo1*^flox/flox^, *CD11c-Cre*, *Piezo1*^-/-^, *Sirt1*^flox/flox^ and *Hif1α*^flox/flox^ mice were obtained from the Jackson Laboratory (Bar Harbor, ME, USA). OT-II TCR-transgenic mice were obtained from the Center of Model Animal Research at Nanjing University (Nanjing, China). CD45.1 mouse was obtained from Beijing University Experimental Animal Center (Beijing, China). C57BL/6 mice were obtained from Fudan University Experimental Animal Center or Beijing University Experimental Animal Center (Beijing, China). All the mice had been backcrossed to the C57BL/6 background for at least eight generations and were used at an age of 6–12-week age old. WT control mice were of the same genetic background and, where relevant, included Cre^+^ mice to account for the effects of Cre (no adverse effects due to Cre expression itself were observed in vitro or in vivo).

### Tumor model

To establish subcutaneous tumors, 5 x 10^5^ MC38 or MC38-OVA tumor cells were injected into C57BL/6 mice, half male and half female, randomization group. These cells formed a tumor of 1 to 2 cm diameter within 2 to 3 weeks of injection and double blinding detection of mouse tumor size.

### Cell isolation from gut-associated lymphatic tissues

Isolation of lamina propria (LP) lymphocytes was performed as described previously(23). The small intestine and large intestine were removed, opened longitudinally, and cut into pieces. After vigorous shaking in HBSS containing EDTA, the supernatants containing epithelial cells and IELs was discarded. The remaining intestinal pieces were digested with collagenase D (Worthington) and pelleted. The pellet was resuspended and placed in a Percoll gradient, and after centrifugation, the interface containing the LP lymphocytes was collected and prepared for further analysis.

### Cell adoptive transfer

Naïve T cells (CD4^+^TCR^+^CD62L^hi^CD44^lo^CD25^-^) from C57BL/6 mice or OT-II TCR-transgenic mice were sorted and transferred into recipient mice. After 24 hrs, the recipient mice were injected s.c. with WT and *Piezo1*^ΔDC^ DCs mixed with OVA_323-339_ in the presence of complete Freund’s adjuvant (CFA; Difco), LPS (Sigma), or 50 kPa hydrogel-conditioned DCs. At Days 8-9 after immunization, DLN cells were harvested and stimulated with their cognate peptides for 2-3 days prior to cytokine mRNA expression and secretion analyses or pulsed with PMA-ionomycin for 5 hrs prior to the intracellular staining of donor-derived T cells.

### Cell Cultures and Flow Cytometry

Spleens were digested with collagenase D, and DCs (CD11c^+^TCR^-^CD19^-^NK1.1^-^F4/80^-^Ly6G^-^) were sorted with a FACSAria II (Becton Dickinson, San Diego, CA, USA). Naïve T cells were sorted from spleen or PLN. For DC-T cell cocultures, DCs and T cells (1:10) were mixed in the presence of 1 μg/ml OVA_323-339_ peptide and 100 ng/ml LPS. After 5 days of culture, live T cells were stimulated with PMA and ionomycin for intracellular cytokine staining or with plate-bound α-CD3 to measure cytokine secretion and mRNA expression. T cell proliferation was determined by pulsing cells with ^3^H-thymidine for the final 12-16 hrs of culture, as described previously(60). For drug treatments, the cells were incubated with vehicle, cyclosporin A (CsA, 10 nM; Sigma), Yoda1 (25 μM, MCE), 11R-VIVIT (100 nM, MCE), ruthenium red (30 μM, Sigma), gadolinium chloride (10 μM, Sigma), 2-deoxy-D-glucose (2-DG, 1 mmol/l, Sigma) or diethyl succinate (1 mmol/l, Sigma) for 0.5-1 hrs before stimulation. For antibody or cytokine treatment, cultures were supplemented with IL-12 (10 μg/ml, Peprotech) and anti-TGFβ1 mAb (20 μg/ml, R&D Systems). Flow cytometry was performed with the following antibodies from eBioscience, BD Biosciences or Abcam: anti-CD11c FITC (N418), anti-CD11c PE (N418), anti-CD11c FITC (N418), anti-CD4 APC-Cy7 (GK1.5; Cat#130-109-536, RRID:AB_2657974), anti-CD8α FITC (53-6.7), anti-CD11b FITC (M1/70), anti-Ly6G PE (RB6-8C5; Cat# ab25378, RRID:AB_470493), anti-F4/80 PE (BM8), anti-CD19 PE (1D3; Cat#340418, RRID:AB_400423), anti-TCR FITC (H57-597), anti-CD44 FITC (IM7), anti-CD62L APC (MEL14), anti-CD80 APC (1C10), anti-CD54 FITC (YN1/1/7.4), anti-MHCII (AF6-120), anti-CD45 APC (30-F11), anti-NK1.1 PE (PK136), anti-IFNγ PE (XMG1.2), anti-IL-4 PE (11B11), anti-Foxp3 PE (FJK-16s), anti-mouse IL-12p40 mAb (241812; Cat#BE0051, RRID:AB_1107698) and anti-mouse TGFβ1 mAb (EPR21143). Flow cytometry data were acquired on a FACSCalibur (Becton Dickinson, CA, USA) or an Epics XL bench-top flow cytometer (Beckman Coulter, CA, USA), and the data were analyzed with FlowJo (RRID:SCR_008520; Tree Star, San Carlos, CA, USA).

### Hydrogel-coated plates

Dow Corning Sylgard 527 (Part A and B, Sigma–Aldrich) was used to prepare PDMS hydrogel-coated plates. Part A and Part B of the gel were mixed to achieve the appropriate tension, as described previously(8, 26, 28). For the 2 kPa gel, the ratio of A:B was 1:2, and for the 50 kPa gel, the ratio of A:B was 0.3. The plates were coated with the hydrogel and incubated overnight at 60 °C. Then, the gels were coated with fibronectin (1 μg/ml, Sigma) for 4 hrs at 37 °C and washed again with PBS.

### Oxygen consumption analysis

Cells were plated in 24-well Seahorse plates at 2 x10^5^ cells per well, and a negative control well containing only media without cells was included. A utility plate containing calibrant solution (1 ml/well) together with the plates containing the injector ports and probes was incubated in a CO_2_-free incubator at 37 °C overnight. The following day, the medium was removed from the cells and replaced with glucose-supplemented XF assay buffer (500 μl/well), and the cell culture plate was incubated in a CO_2_-free incubator for at least 0.5 hrs. Inhibitors (oligomycin, carbonyl cyanide-4-[trifluoromethoy]) phenylhydrazone, 2-DG, and rotenone (70 μl) were added to the appropriate port of the injector plate. This plate, together with the utility plate, was run on the Seahorse for calibration. Then, the utility plate was replaced with the cell culture plate, and the cell culture plate was analyzed on the Seahorse XF-24 instrument.

### Measurement of intracellular Ca^2+^ concentrations

The intracellular Ca^2+^ concentrations ([Ca^2+^]) were measured fluorometrically using the fluorescent calcium indicator dye Fura2 AM (Sigma), as previously described(25). Cells were incubated with 5 μM Fura2 AM in HBSS supplemented with 110 mM NaCl, 5 mM KCl, 0.3 mM Na_2_HPO_4_, 0.4 M KH_2_PO_4_, 5.6 mM glucose, 0.8 mM MgSO_4_, 7 mM H_2_O, 4 mM NaHCO_3_, 1.26 mM CaCl_2_, and 15 mM HEPE, at pH 7.4, at room temperature for 60 min.

### RNA and protein expression analysis

RNA was extracted with a RNeasy kit (QIAGEN, Dusseldorf, Germany), and cDNA was synthesized using SuperScript III reverse transcriptase (Invitrogen, Carlsbad, CA). An ABI 7900 real-time PCR system was used for quantitative PCR, with primer and probe sets obtained from Applied Biosystems (Carlsbad, CA). The results were analyzed using SDS 2.1 software (Applied Biosystems). The cycling threshold value of the endogenous control gene (*Hprt*1, which encodes hypoxanthine guanine phosphoribosyl transferase) was subtracted from the cycling threshold (ΔC_T_). The expression of each target gene is presented as the fold change relative to that of control samples (2-^ΔΔCT^). For the detection of phosphorylated signaling proteins, purified cells were activated with LPS (Sigma), immediately fixed with Phosflow Perm buffer (BD Biosciences) and stained with phycoerythrin or allophycocyanin directly conjugated to antibodies against Erk phosphorylated at Thr202 and Tyr204 (20A; Cat#612566, RRID:AB_399857; BD Biosciences), p38MAPK phosphorylated at Thr180 and Thr182 (D3F9; Cell Signaling Technology), JNK phosphorylated at Thr183 and Tyr185 (G9; Cell Signaling Technology), STAT4 phosphorylated at Tyr701 and Ser727 (58D6; Cell Signaling Technology), and SMAD3 phosphorylated at Tyr705 and Ser727 (D3A7; Cell Signaling Technology), as described previously(60). Intracellular staining analysis was performed as described(23) using anti-HIF-1α (EPR16897; Abcam) and anti-SIRT1 (17A7AB4; Abcam) antibodies.

### IL-12Rβ2 and TGFβR2 knockdown with RNAi

A gene-knockdown lentiviral construct was generated by subcloning gene-specific short hairpin RNA (shRNA) sequences into lentiviral shRNA expression plasmids (pMagic4.1) as described previously(50). Lentiviruses were harvested from the culture supernatant of 293T cells (KCB Cat# KCB 200744YJ, RRID:CVCL_0063) transfected with shRNA vector. Sorted OT-II CD4^+^ T cells were infected with the recombinant lentivirus, and green fluorescent protein-expressing cells were isolated using fluorescence sorting 48 hrs later. IL-12Rβ2 and TGFβR2 expression was confirmed using real-time PCR. The sorted T cells expressing either control or shRNA vectors were used for functional assays.

### Statistical analysis

All the data are presented as the mean ± SD. Student’s unpaired *t* test was used for the comparison of means to evaluate differences between groups. Analysis of the survival curves was performed using the log-rank (Mantel–Cox) test. A *P* value (alpha-value) of less than 0.05 was considered statistically significant.

## Appendix 1

### Appendix 1-key resources table

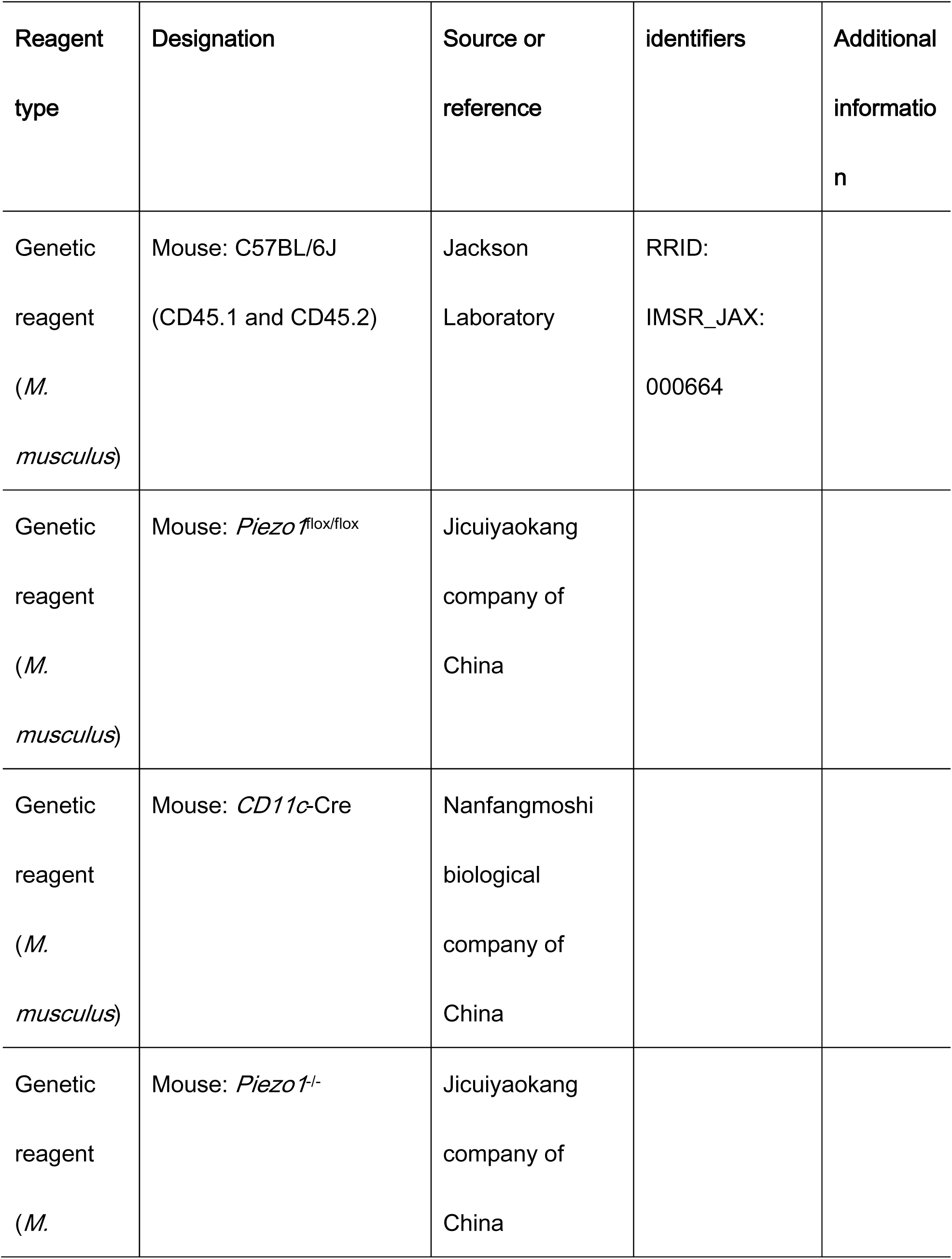

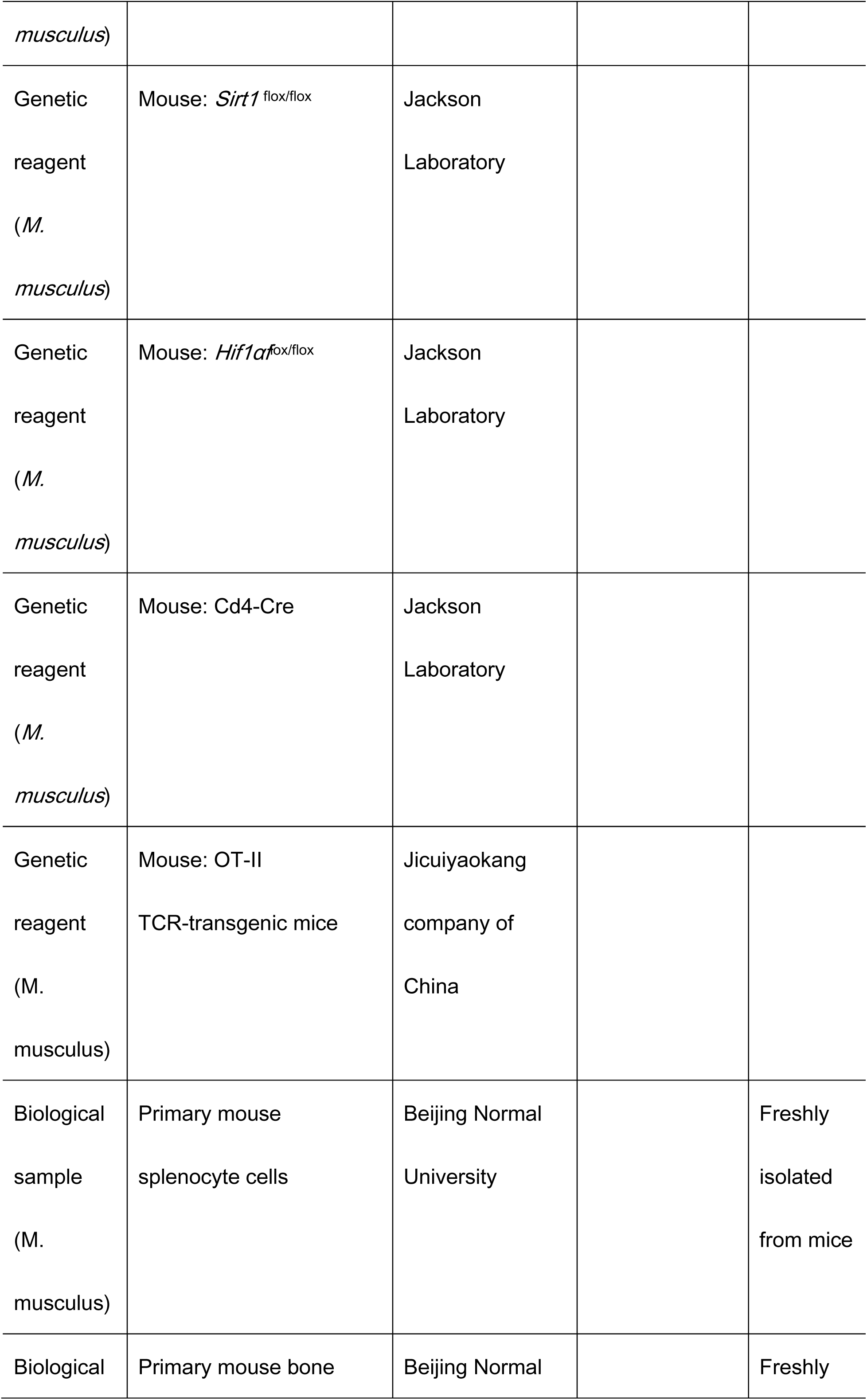

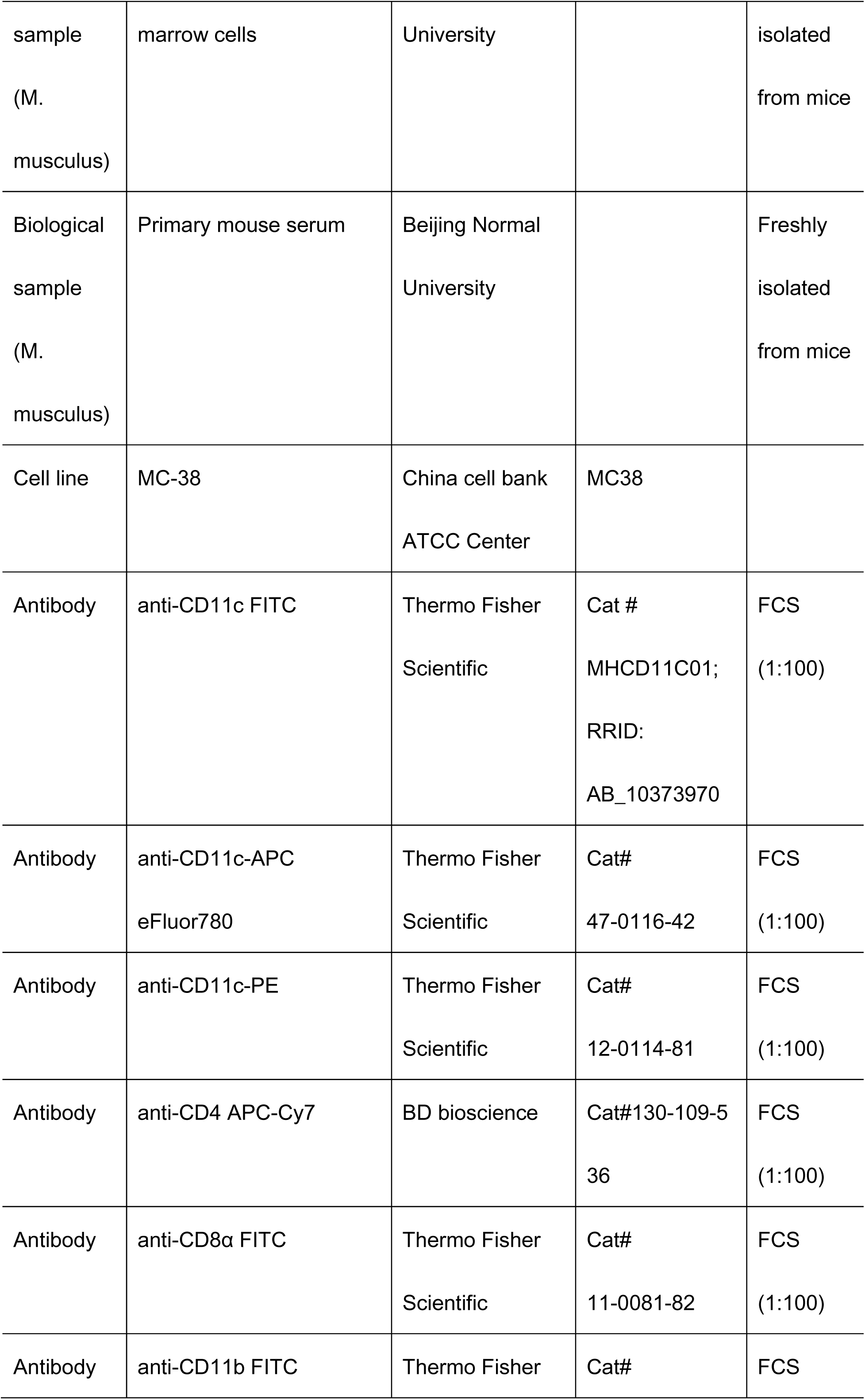

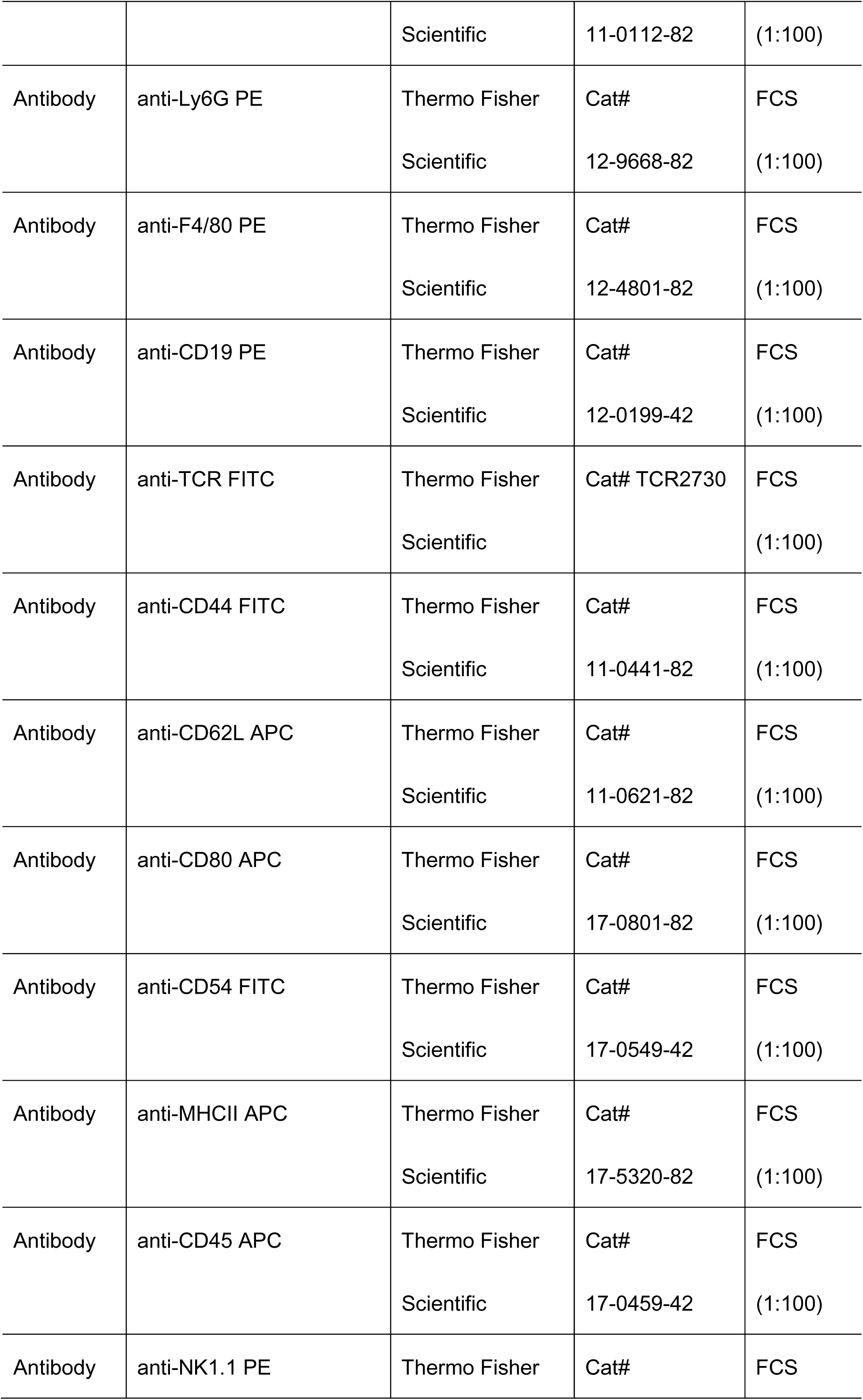

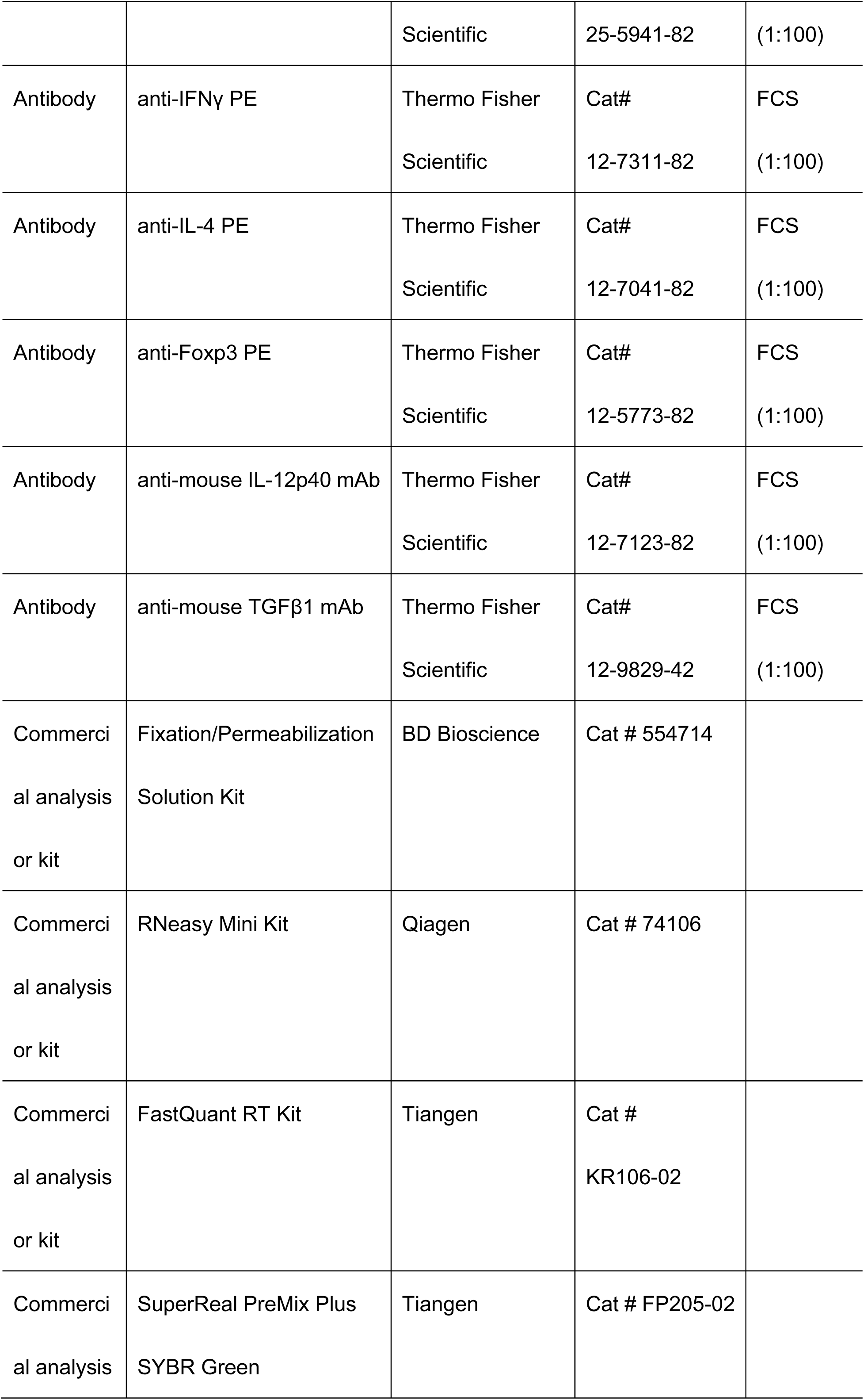

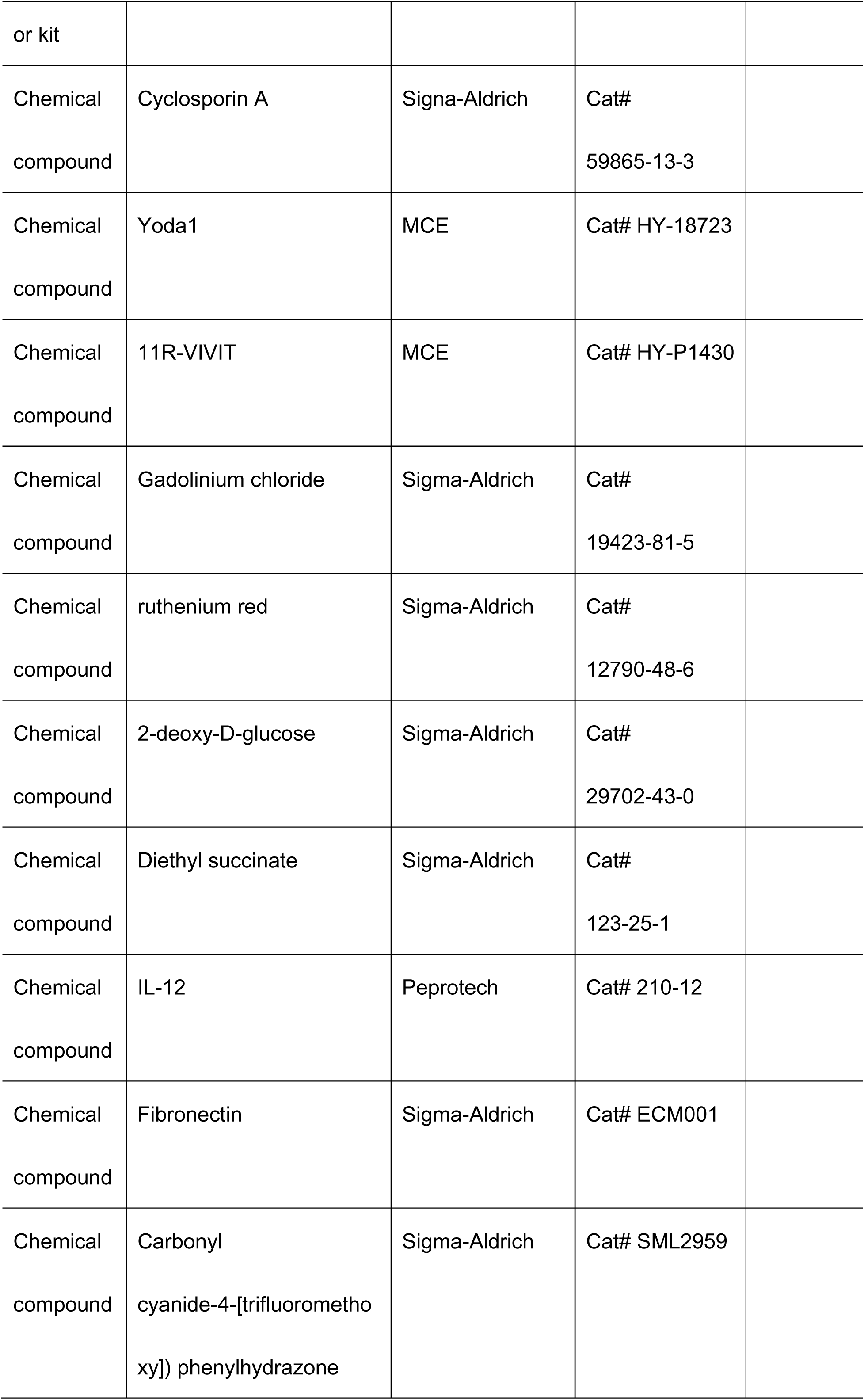

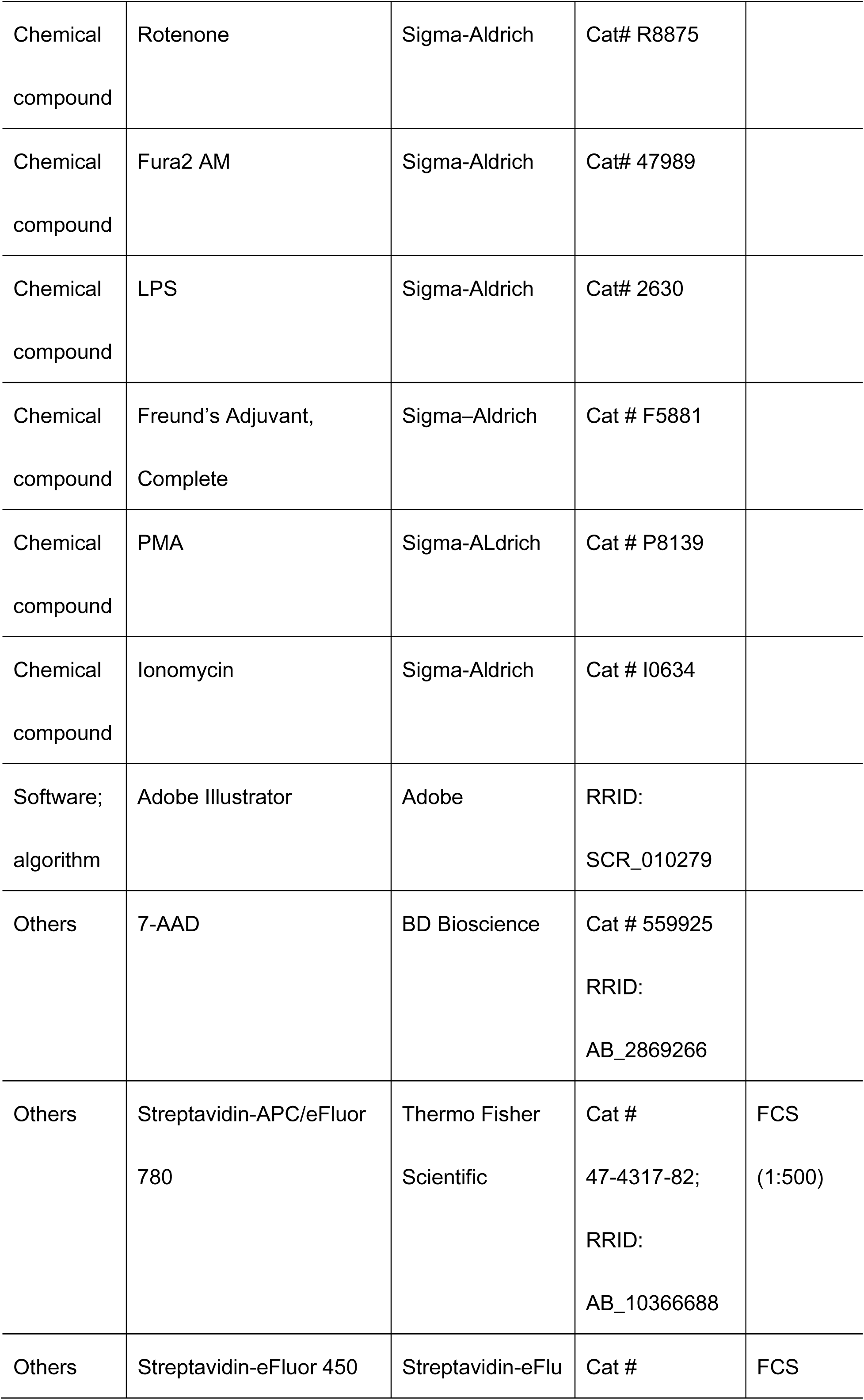

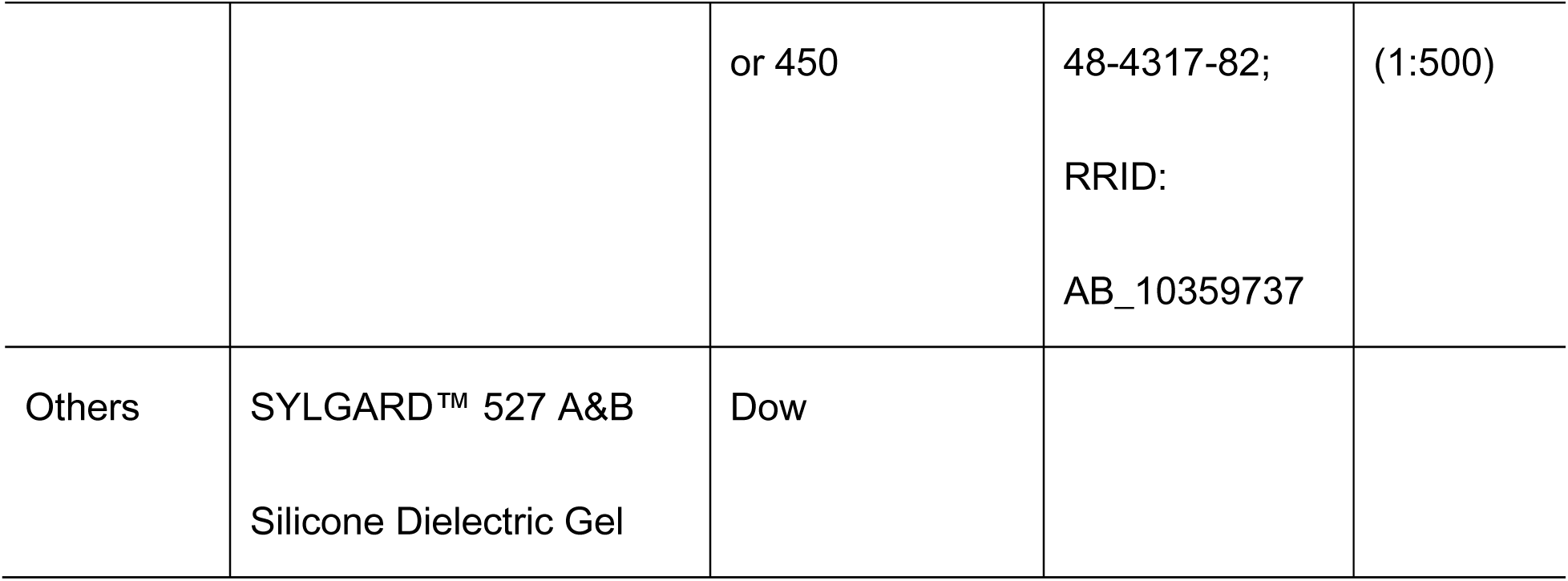

## Acknowledgements

The authors’ research is supported by grants from the National Natural Science Foundation for Key Programs of China (31730024, G.L.), National Natural Science Foundation for General Programs of China (32170911, G.L. and 31970863, Y.B.) and Beijing Municipal Natural Science Foundation of China (5202013, G.L.).

## Author contributions

Y.X.W. conducted the experiments with cells and mice and contributed to the mouse colonies management; H.Y. designed and conducted the experiments with mice and contributed to the mouse colonies management; A.J. conducted the experiments with cells and analyzed data; Y.F.W. contributed to the experiments with mice and contributed to the mouse colonies management; Q.Y. and Y.D., contributed to the discussion of overall studies; Y.B. designed and conducted the experiments with cells and mice, analyzed data and contributed to the manuscript writing; G.L. developed the concept, designed, and conducted the experiments with cells and mice, analyzed data, wrote the manuscript, and provided overall direction.

## COMPETING FINANCIAL INTERESTS

The authors declare no competing financial interests.

---

**Supplementary Figure 1.**
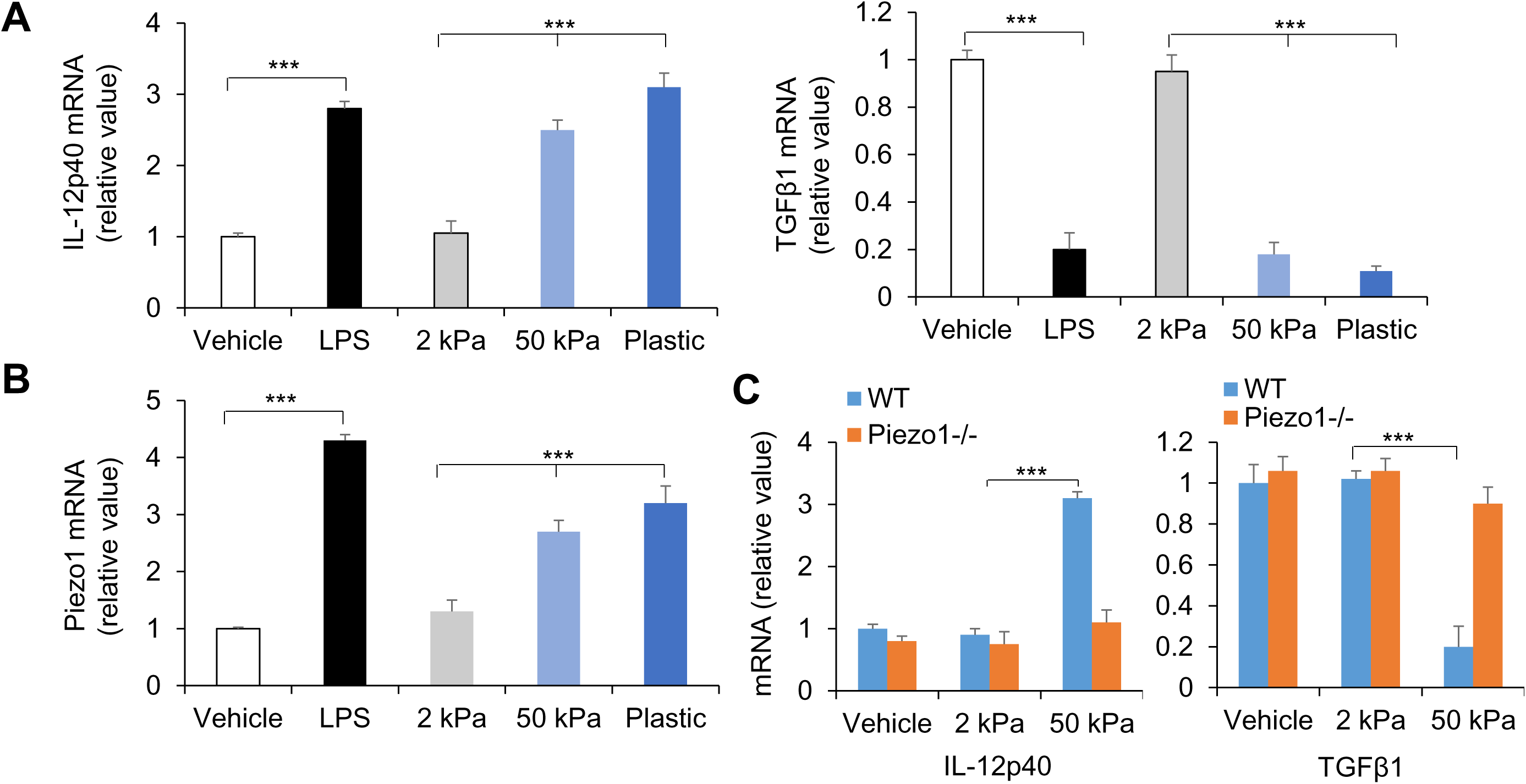
Piezo1 expressions of DCs following innate stimuli. IL-12p40 and TGFβ1 mRNA (**A & C**) and Piezo1 mRNA (**B**) expression of splenic DCs from C57BL/6 or/and *Piezo1*^-/-^ mice with the indicated treatment (LPS, 10 ng/mL or conditioned with 2-kPa or 50 kPa-hydrogels plate). Levels in the vehicle group were set to 1. Data are representative three to four independent experiments (mean ± s.d.; n=3-4). \*\*\**P*<0.001, compared with the indicated groups.

**Supplementary Figure 2.**
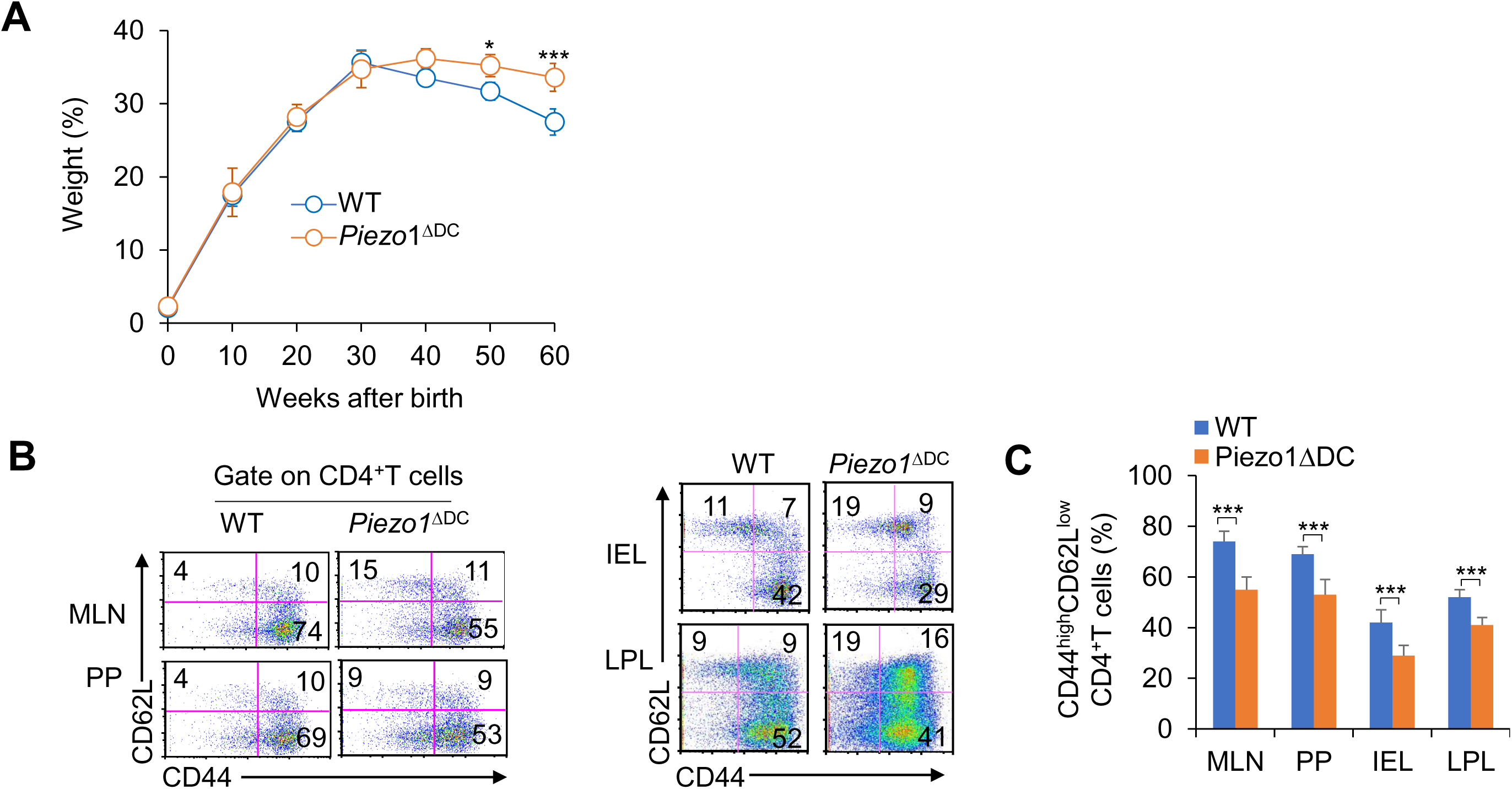
DC-specific Piezo1 deficiency reduced T cell activities in aged mice. (**A**) Body weights of WT and *Piezo1*^ΔDC^ mice are shown (n=20). (**B**) Piezo1 deficiency in DCs resulted in more CD44^high^CD62^Low^ cells in the CD4^+^ T cell population than WT DCs. A typical figure shown in **B** and data summarized in **C**. The data are representative of three independent experiments (mean ± s.d.; n=4-20). \**P*<0.05 and \*\*\**P*<0.001, compared with the indicated groups.

**Supplementary Figure 3.**
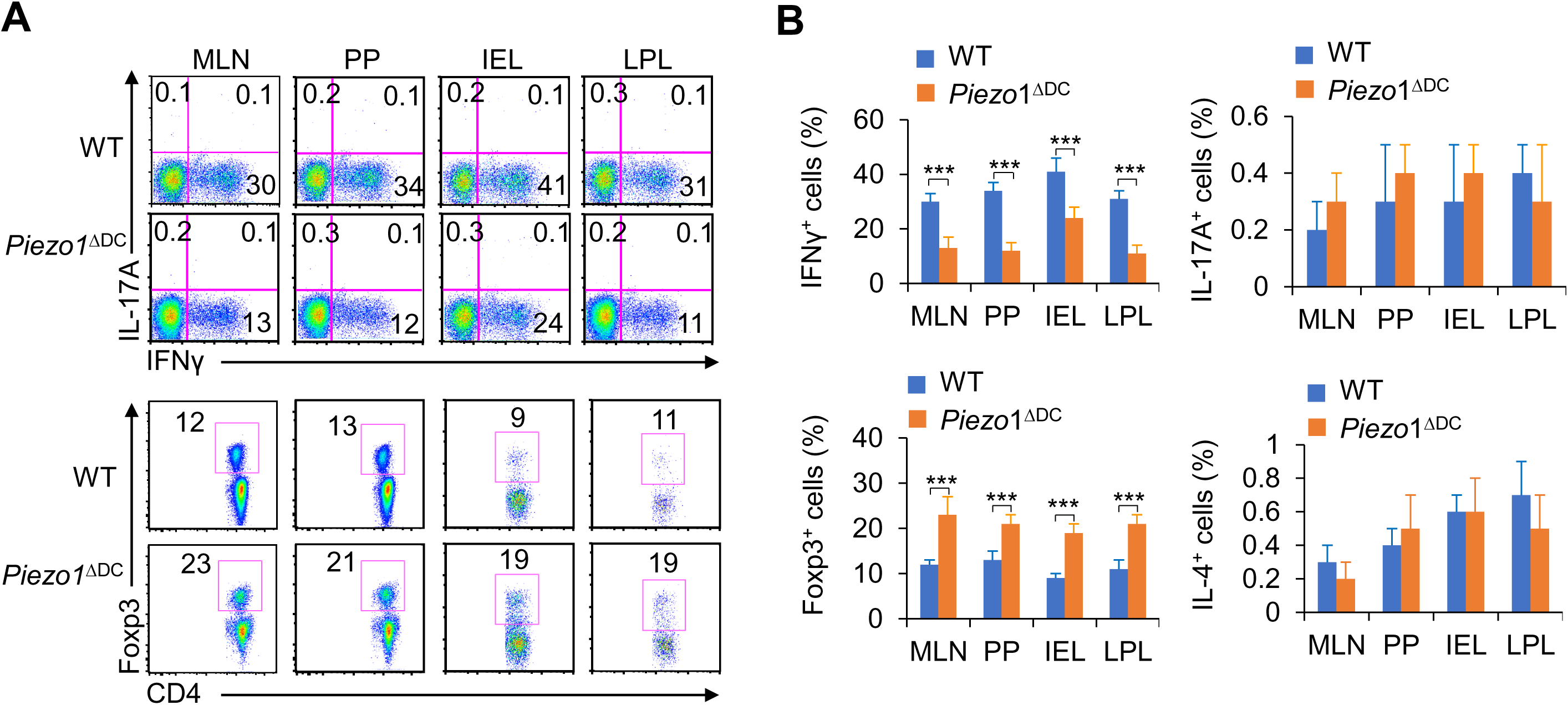
DC-specific Piezo1 deficiency alters T cell differentiation in the aged mice. Intracellular IFNγ, IL-4, IL-17A and Foxp3 expression by CD4^+^ T cells from the MLN, PP, IEL and LPL in WT and *Piezo1*^ΔDC^ mice. A representative figure is shown in (**A**), and the percentage of positive cells is shown in (**B**). The data are representative of three to four independent experiments (mean ± s.d.; n=4-5). \*\*\**P*<0.001, compared with the indicated groups.

**Supplementary Figure 4.**
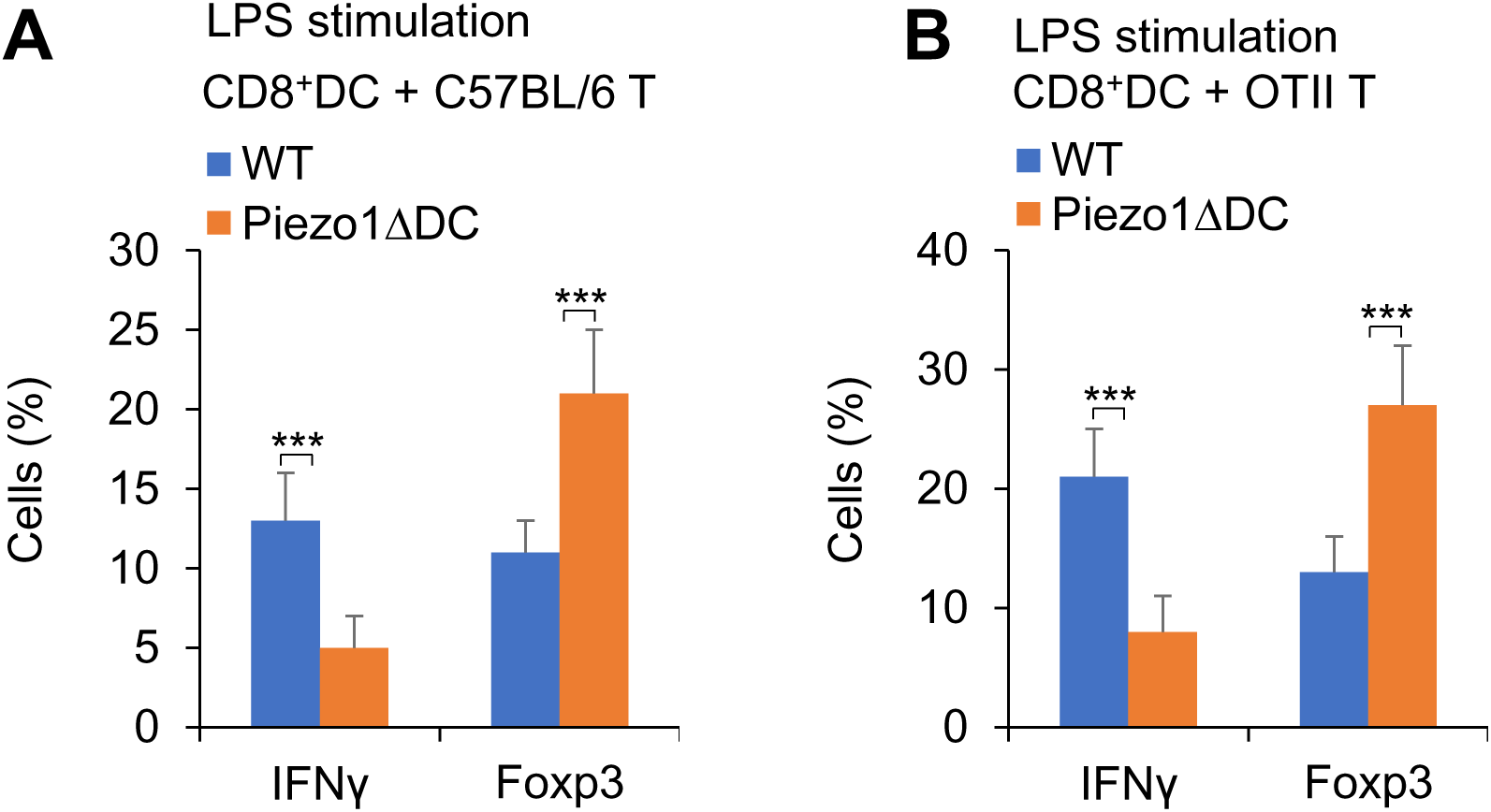
DC Piezo1 directs T_H_1 and T_reg_ differentiation in vitro. (**A**) C57BL/6 naïve CD4^+^T cells cocultured with LPS-pulsed CD8^+^ DCs from WT and *Piezo1*^ΔDC^ mice for 5 days. Intracellular staining of IFNγ and Foxp3 among T cells. (**B**) Naïve CD4^+^ T cells from OT II mice cocultured with LPS-pulsed CD8^+^ DCs from WT and *Piezo1*^ΔDC^ mice for 5 days. Intracellular staining of IFNγ and Foxp3 among T cells. Data are representative three to four independent experiments (mean ± s.d.; n=3-5). \*\*\**P*<0.001, compared with the indicated groups.

**Supplementary Figure 5.**
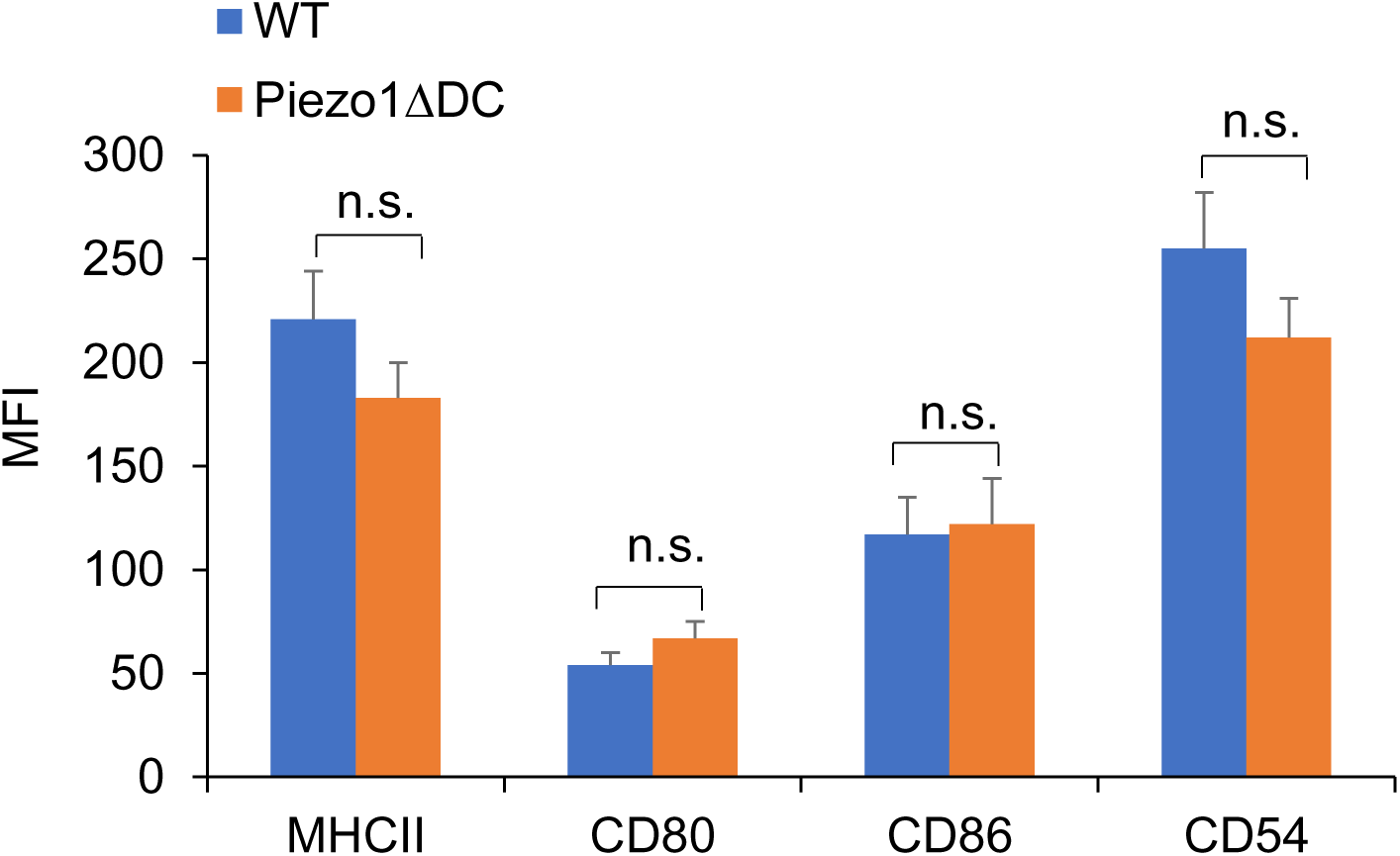
Piezo1 regulates DC IL-12 and TGFβ1 production in directing T_H_1 and T_reg_ cell differentiation. Expression of indicated molecules in WT and *Piezo1*^ΔDC^ splenic DCs in the presence of LPS (10 ng/mL). Data are representative three to four independent experiments (mean ± s.d.; n=3). n.s., not significant.

**Supplementary Figure 6.**
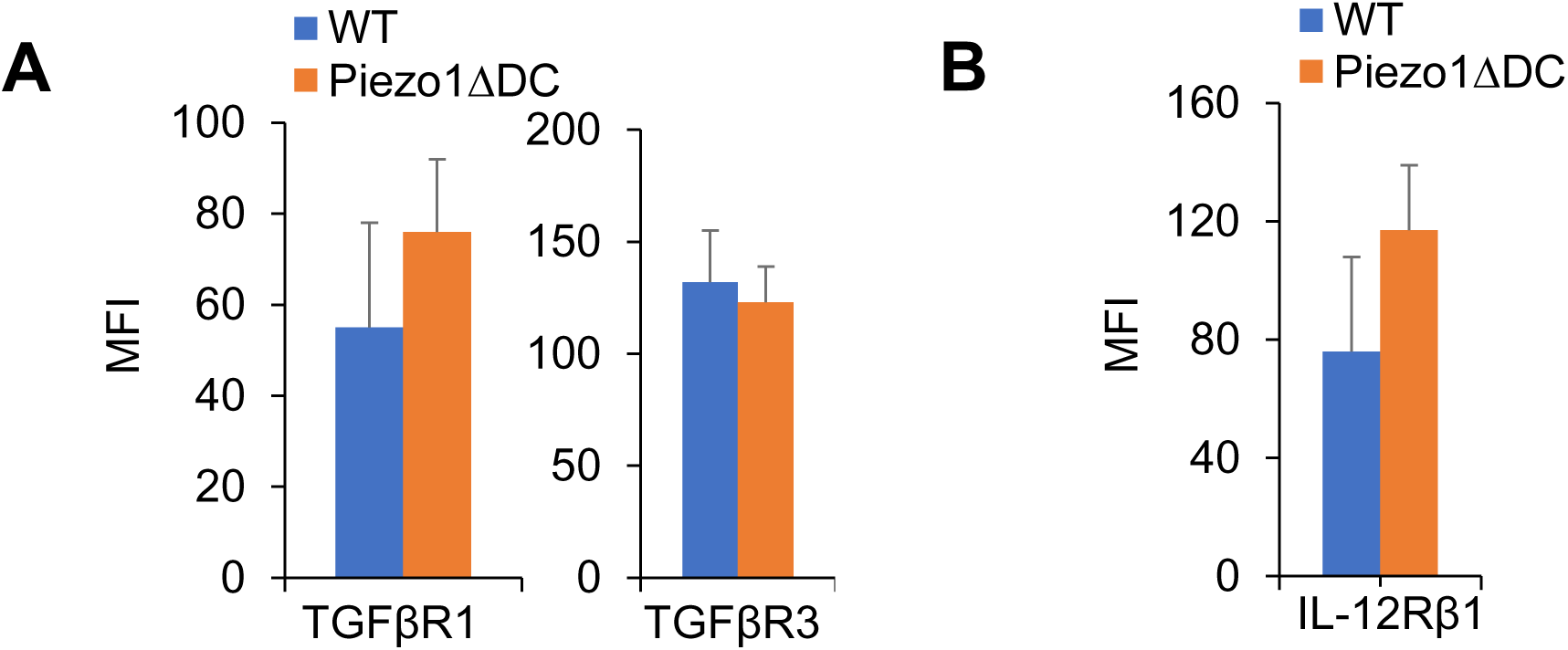
T cell TGFβR2-Smad3 and IL-12Rβ2-pSTAT4 are necessary for DC Piezo1 signaling. Expression of TGFβR1, TGFβR3 (**A**) and IL-12Rβ1 (**B**) in T cells cocultured with WT or *Piezo1*^ΔDC^ splenic DCs for 5 days. Data summarized. Data are representative three to four independent experiments (mean ± s.d.; n=4).

**Supplementary Figure 7.**
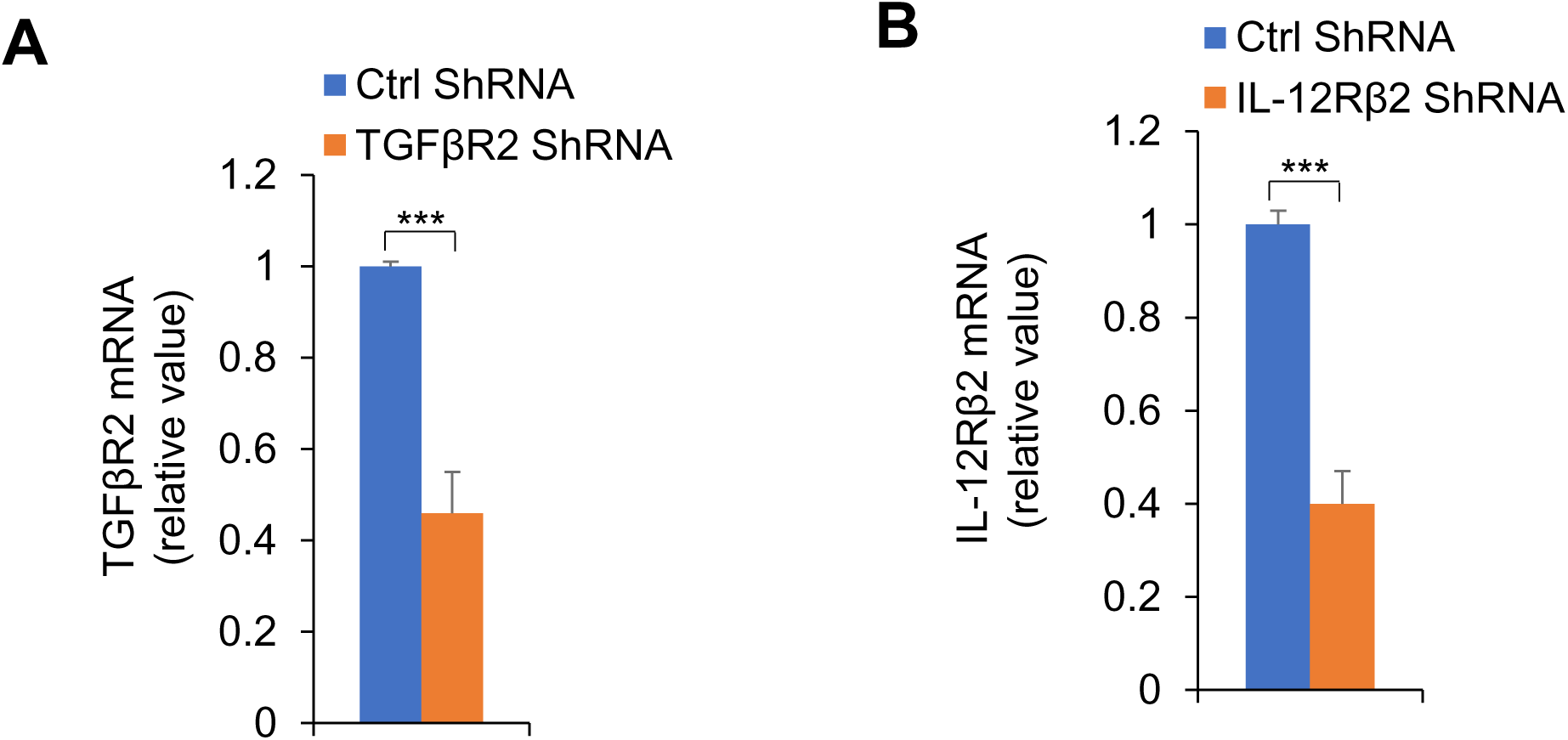
T cell TGFβR2-Smad3 and IL-12Rβ2-pSTAT4 are necessary for DC Piezo1 signaling. Sorted CD4^+^ T cells were transfected with control or TGFβR2 shRNA vector (**A**) or IL-12Rβ2 shRNA vector (**B**) and stimulated with WT or *Piezo1*^ΔDC^ splenic DCs for 5 days in the presence of LPS (10 ng/ml). Expression of indicated mRNA (A or B) were determined with qPCR (Levels of ctrl siRNA were set to 1). Data are representative of three independent experiments (mean ± s.d.; n=3). ***P<0.001 compared with the indicated groups.

**Supplementary Figure 8.**
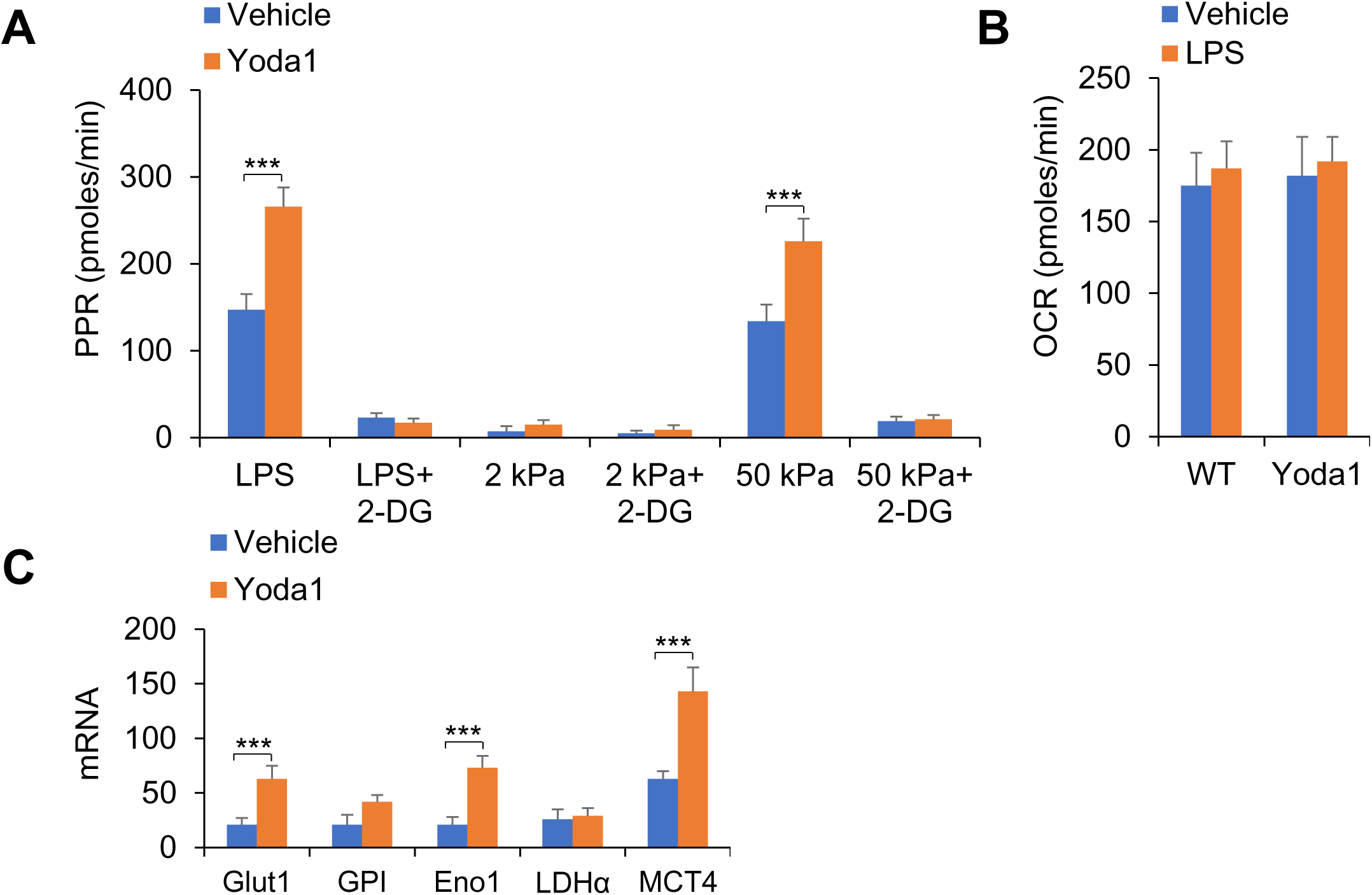
Piezo1 regulates TGFβ1 and IL-12 production through SIRT1-HIF1α-glycolysis pathway. Sorted splenic DCs from WT mice were stimulated with LPS (10 ng ml^-1^) or with 2 kPa or 50 kPa-hydrogels conditions for 24 hrs in the presence or absence of 2-DG (1 mmol) or Yoda1 (25 μM). (**A**) The PPR was analyzed as a readout for glycolysis. (**B**) The OCR was analyzed as a readout for oxidative phosphorylation (OXPHOS). (**C**) mRNA expression of glycolytic molecules in splenic DCs from WT mice in the presence of LPS (10 ng/mL) and Yoda1 (25 μM). Levels in the WT control groups were set to 1. Data are representative three to four independent experiments (mean ± s.d.; n=4). \**P*<0.05, \*\**P*<0.01 and \*\*\**P*<0.001, compared with the indicated groups.

**Supplementary Figure 9.**
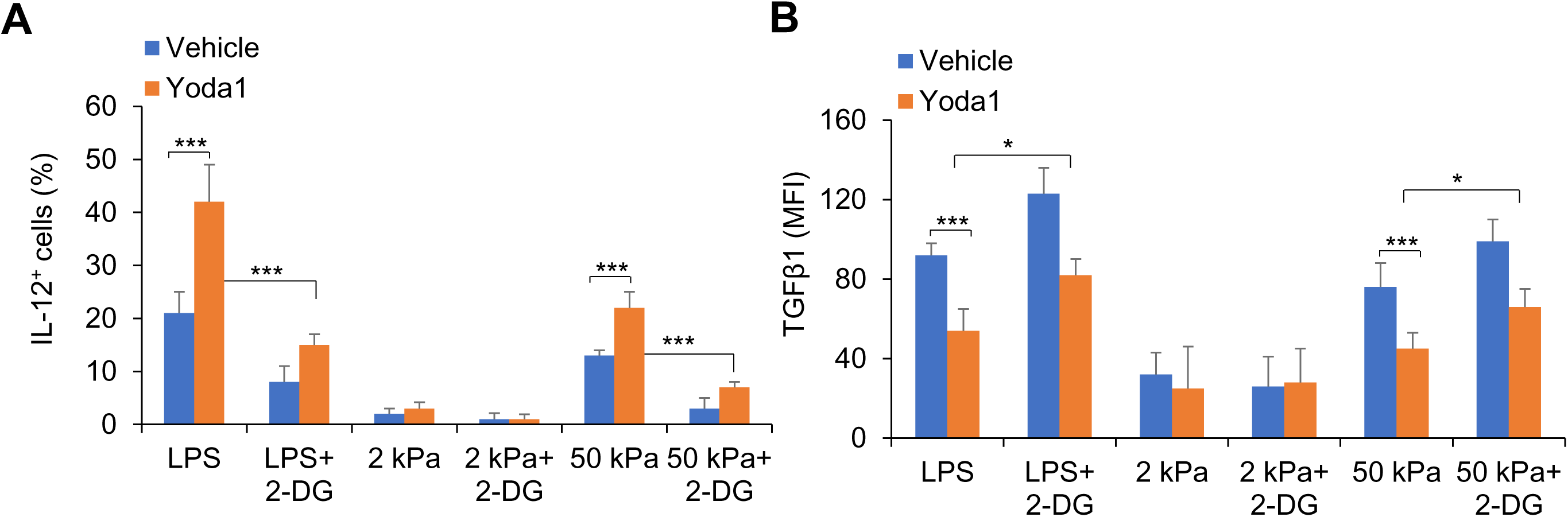
Piezo1 regulates TGFβ1 and IL-12 production through SIRT1-HIF1α-glycolysis pathway. (**A-B**) Sorted splenic DCs from WT mice were stimulated with LPS (10 ng ml^-1^) or with 2 kPa or 50 kPa-hydrogels conditions for 24 hrs in the presence or absence of 2-DG (1 mmol) or Yoda1 (25 μM). Intracellular staining of IL-12p40 (**A**) and TGFβ1 (**B**) expression with indicated treatment. Data are representative three to four independent experiments (mean ± s.d.; n=3). \**P*<0.05 and \*\*\**P*<0.001, compared with the indicated groups.

**Supplementary Figure 10.**
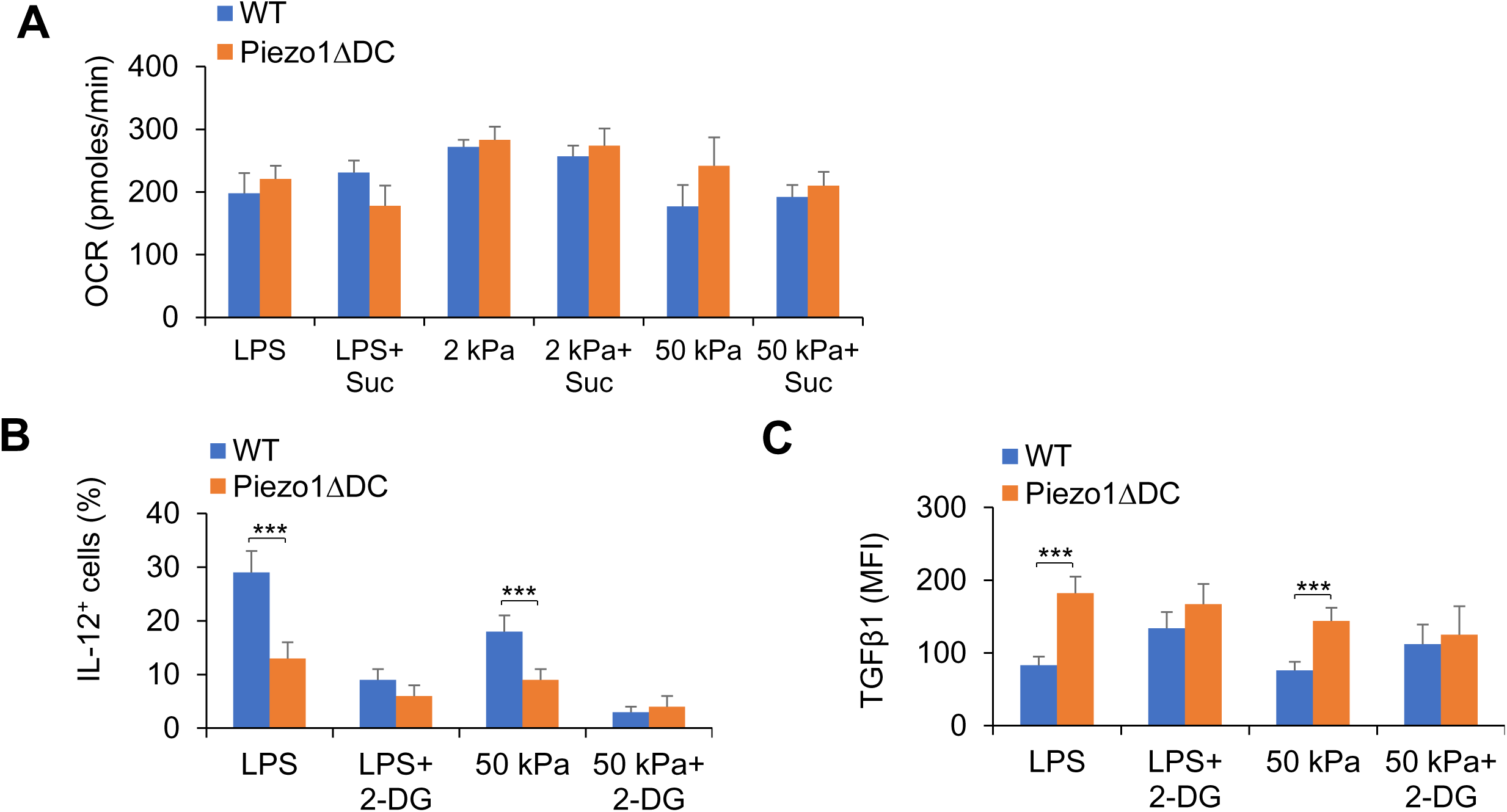
Piezo1 regulates TGFβ1 and IL-12 production through SIRT1-HIF1α-glycolysis pathway. (**A**) Sorted splenic DCs from WT or *Piezo1*^ΔDC^ mice were stimulated with LPS (10 ng/ml) or 2 kPa or 50 kPa hydrogels for 24 hrs in the presence or absence of succinate (5 mmol/l). The OCR was analyzed as a readout for OXPHOS. (**B-C**) Sorted splenic DCs from WT or *Piezo1*^ΔDC^ mice were stimulated with LPS (10 ng ml-1) or with 2 kPa or 50 kPa-hydrogels conditions for 24 hrs in the presence or absence of 2-DG (1 mmol). Intracellular staining of IL-12p40 (B) and TGFβ1 (C) expression with indicated treatment. Data are representative three independent experiments (mean ± s.d.; n=3). \*\*\**P*<0.001, compared with the indicated groups.

**Supplementary Figure 11.**
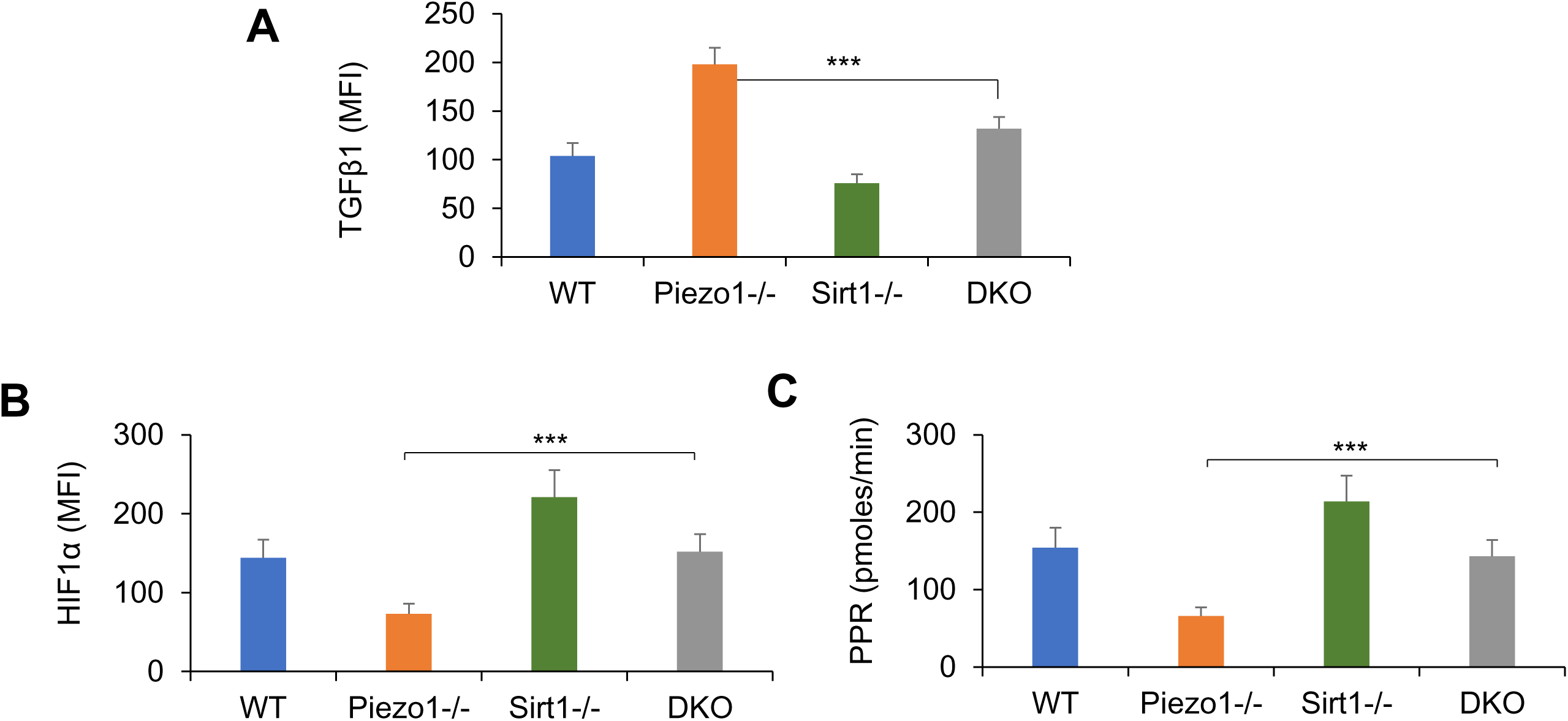
Piezo1 regulates TGFβ1 and IL-12 production through SIRT1-HIF1α-glycolysis pathway. Splenic DCs from WT, *Piezo1*^ΔDC^ (*Piezo1*^-/-^), *Sirt1*^-/-^, and Piezo1/Sirt1 double knockout (DKO) mice were stimulated with LPS (10 ng/mL). (**A**) Intracellular staining of TGFβ1. (**B**) Intracellular staining of HIF1α. (**C**) PPR was analyzed as a readout for glycolysis. Data summarized. Data are representative three to four independent experiments (mean ± s.d.; n=3-4). \*\*\**P*<0.001, compared with the indicated groups.

**Supplementary Figure 12.**
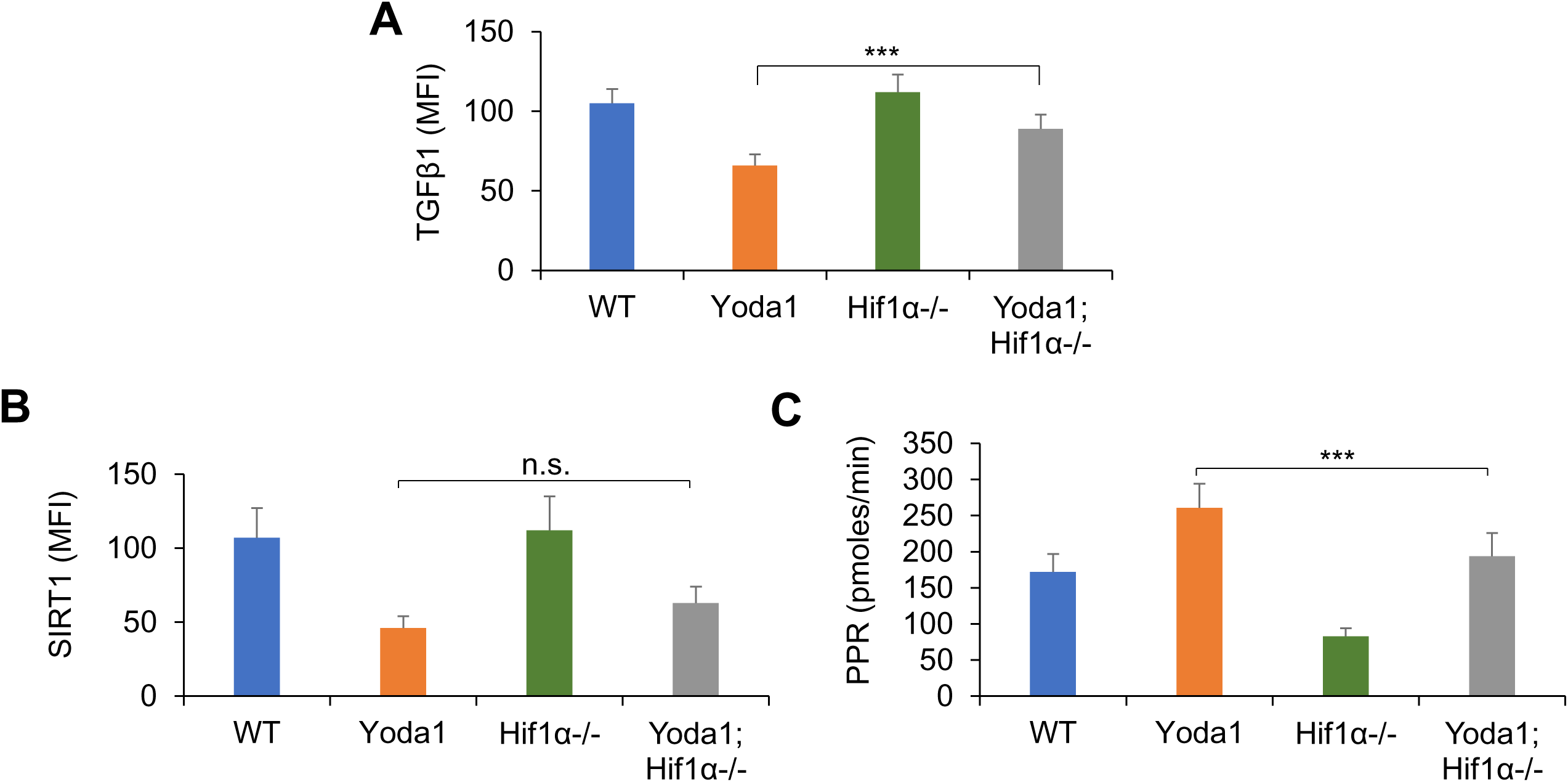
Piezo1 regulates TGFβ1 and IL-12 production through SIRT1-HIF1α-glycolysis pathway. Splenic DCs from WT or *Hif1α*^-/-^ mice were stimulated with LPS (10 ng/mL) in the presence or absence of Yoda1 (25 μM). (**A**) Intracellular staining of TGFβ1. (**B**) Intracellular staining of SIRT1. (**C**) PPR was analyzed as a readout for glycolysis. Data are representative three to four independent experiments (mean ± s.d.; n=3-4). \*\*\**P*<0.001, compared with the indicated groups.

**Supplementary Figure 13.**
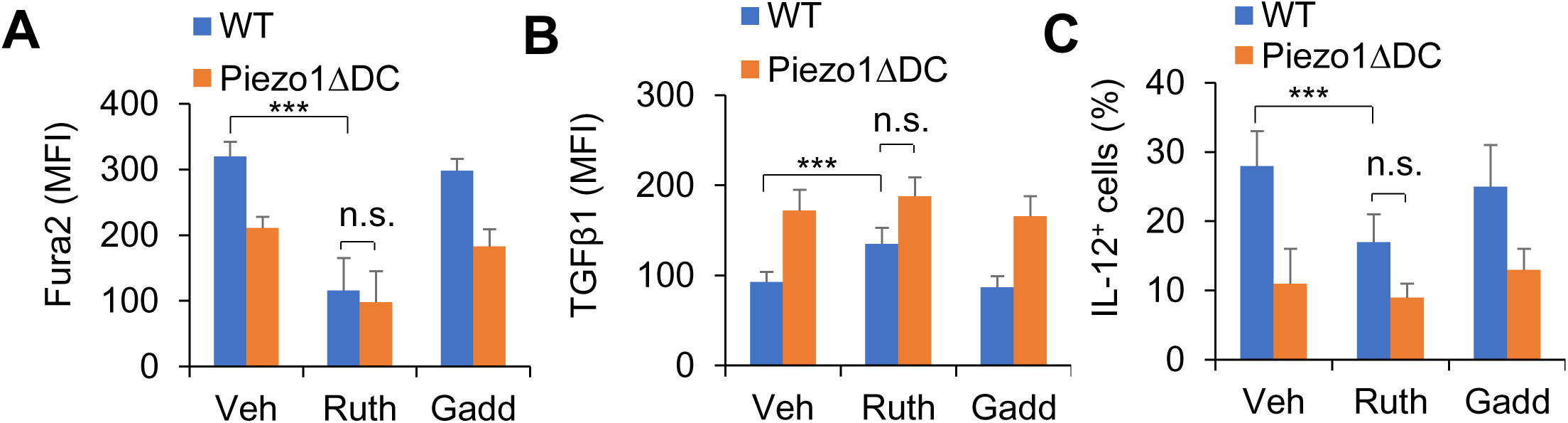
Piezo1 regulates TGFβ1 and IL-12 production through Ca2^+^- Calcineurin-NFAT axis. (**A**) Intracellular Ca^2+^ using Fura2 in splenic DCs from WT or *Piezo1*^ΔDC^ mice with indicated treatment (Yoda1, 25 μM; Ruthenium red, 30 μM, Sigma; Gadolinium chloride, 10 μM, Sigma). (**B-C**) Intracellular staining of TGFβ1 (B) and IL-12p40 (C) in splenic DCs from WT or *Piezo1*^ΔDC^ mice with indicated treatments. Data are representative three to four independent experiments (mean ± s.d.; n=3). \*\*\**P*<0.001, compared with the indicated groups. n.s., not significant.

**Supplementary Figure 14.**
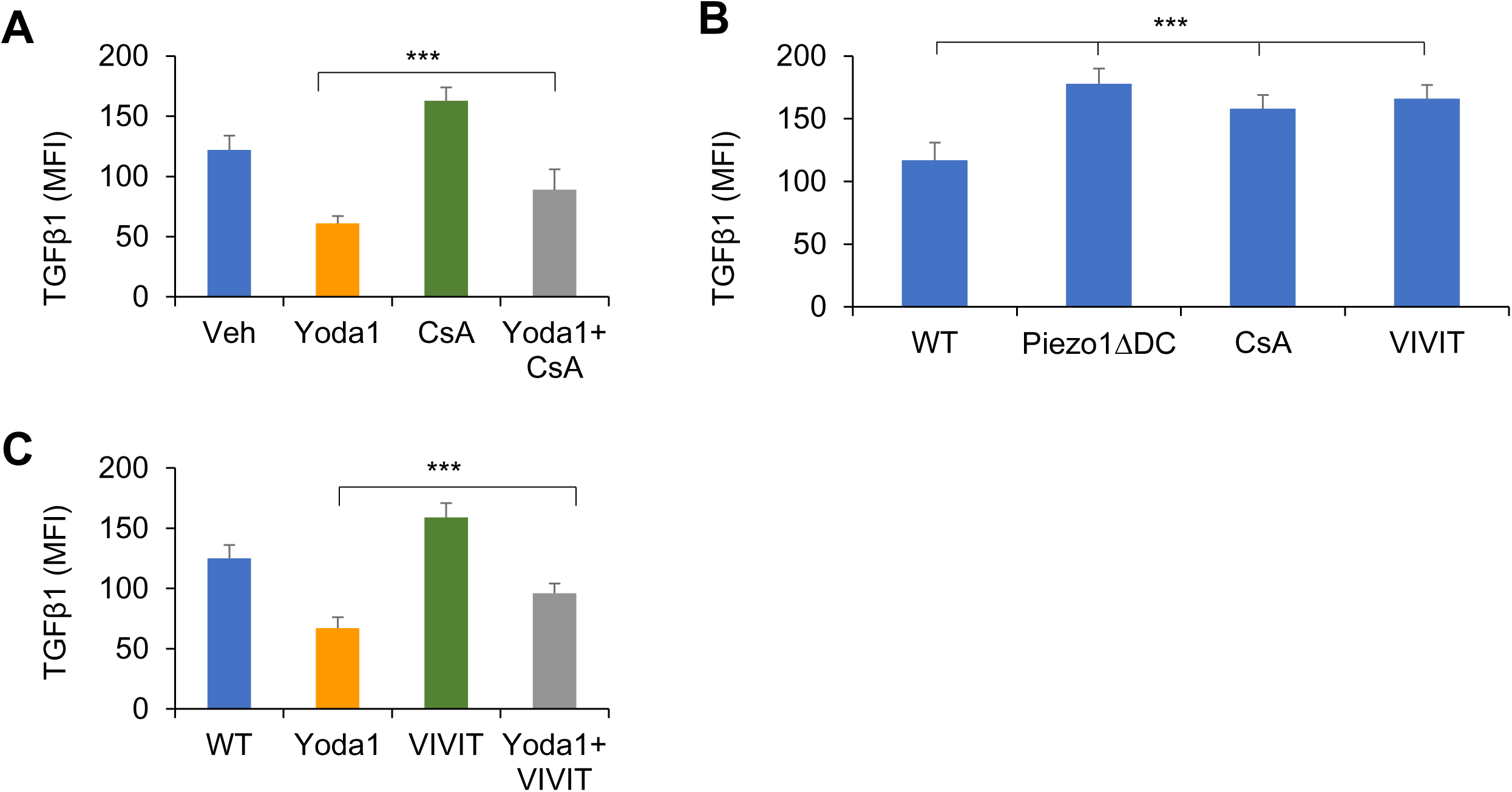
Piezo1 regulates TGFβ1 and IL-12 production through Ca2+- Calcineurin-NFAT axis. (**A or C**) Intracellular staining of TGFβ1 in splenic DCs from WT mice with indicated treatments. (**B**) Intracellular staining of TGFβ1 in splenic DCs from WT or *Piezo1*^ΔDC^ mice with indicated treatments (Yoda1, 25 μM, MCE; 11R-VIVIT, 100 nM, MCE; CsA, 10 nM, Sigma). Data are representative three to four independent experiments (mean ± s.d.; n=3-5). \*\*\**P*<0.001, compared with the indicated groups.

**Supplementary Figure 15.**
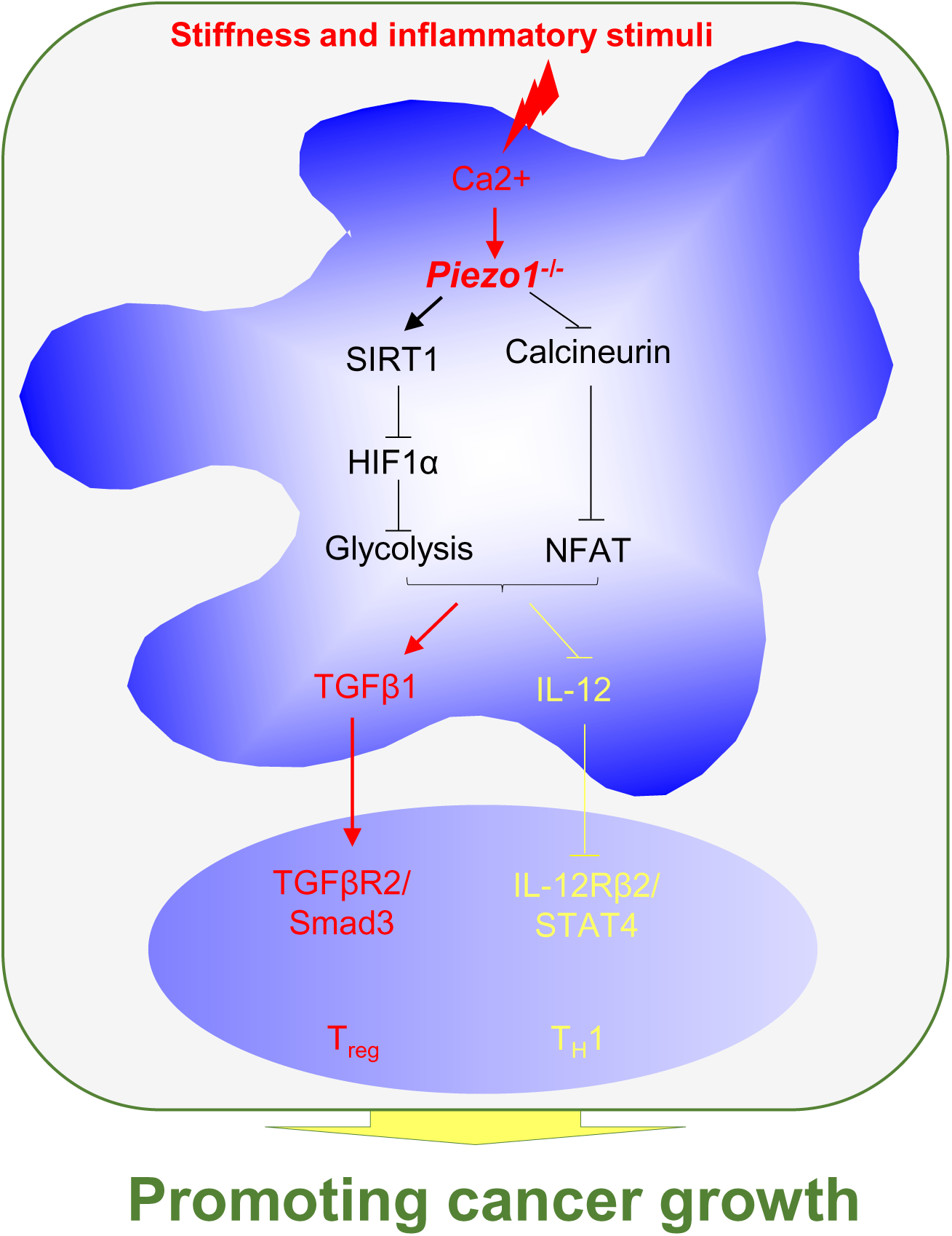
DC Piezo1 control the differentiation of T_H_1 ad T_reg_ cells in cancer. Proposed model of how Piezo1 in DCs integrates inflammatory stimuli and stiffness to regulate the differentiation of T_H_1 and T_reg_ cell populations in promoting cancer growth.

## Notes

### Competing Interest Statement

The authors have declared no competing interest.

